# Mg^2+^-dependent conformational equilibria in CorA: an integrated view on transport regulation

**DOI:** 10.1101/2021.08.20.457080

**Authors:** Nicolai Tidemand Johansen, Marta Bonaccorsi, Tone Bengtsen, Andreas Haahr Larsen, Frederik Grønbæk Tidemand, Martin Cramer Pedersen, Pie Huda, Jens Berndtsson, Tamim Darwish, Nageshwar Rao Yepuri, Anne Martel, Thomas Günther Pomorski, Andrea Bertarello, Mark Sansom, Mikaela Rapp, Ramon Crehuet, Tobias Schubeis, Kresten Lindorff-Larsen, Guido Pintacuda, Lise Arleth

**Affiliations:** Condensed Matter Physics, Niels Bohr Institute, University of Copenhagen, Copenhagen, Denmark; Centre de RMN à Très hauts Champs de Lyon (UMR 5280, CNRS / Ecole Normale Supérieure de Lyon / Université Claude Bernard Lyon 1), University of Lyon, Villeurbanne, France; Structural Biology and NMR Laboratory and Linderstrøm-Lang Centre for Protein Science, Department of Biology, University of Copenhagen, Denmark; Department of Biochemistry, University of Oxford, Oxford, United Kingdom; Australian Institute for Bioengineering and Nanotechnology, The University of Queensland, Brisbane, Australia; Department of Biochemistry and Biophysics, Center for Biomembrane Research, Stockholm University, Stockholm, Sweden; National Deuteration Facility, Australian Nuclear Science and Technology Organization, Lucas Heights, Australia; Institut Laue–Langevin, Grenoble, France; Section for Transport Biology, Department of Plant and Environmental Sciences, University of Copenhagen, Frederiksberg, Denmark; Department of Biochemistry II – Molecular Biochemistry, Ruhr University Bochum, Bochum, Germany

## Abstract

The CorA family of proteins regulates the homeostasis of divalent metal ions in many bacteria, archaea, and eukaryotic mitochondria, making it an important target in the investigation of the mechanisms of transport and its functional regulation. Although numerous structures of open and closed channels are now available for the CorA family, the mechanism of the transport regulation remains elusive. Here, we investigated the conformational distribution and associated dynamic behaviour of the pentameric Mg^2+^ channel CorA at room temperature using small-angle neutron scattering (SANS) in combination with molecular dynamics (MD) simulations and solid-state nuclear magnetic resonance spectroscopy (NMR). We find that neither the Mg^2+^-bound closed structure nor the Mg^2+^-free open forms are sufficient to explain the average conformation of CorA. Our data support the presence of conformational equilibria between multiple states, and we further find a variation in the behaviour of the backbone dynamics with and without Mg^2+^. We propose that CorA must be in a dynamic equilibrium between different non-conducting states, both symmetric and asymmetric, regardless of bound Mg^2+^ but that conducting states become more populated in Mg^2+^-free conditions. These properties are regulated by backbone dynamics and are key to understanding the functional regulation of CorA.

## Introduction

Magnesium is the most abundant divalent cation (Mg^2+^) inside the cell, where it is mainly associated with the biological energy source adenosine triphosphate and other negatively charged molecules^1^. Mg^2+^ serves several biological functions, e.g. as co-factor for enzymes^1^, and Mg^2+^ deficiency is linked to severe diseases including cardiac syndromes, muscular dysfunction and bone wasting^2–4^. CorA is the main ion channel for Mg^2+^-import in most bacteria and archea^5^. Despite little sequence conservation, CorA shares two membrane spanning helices and a conserved GMN motif with eukaryotic homologs, including Mrs2 that is responsible for Mg^2+^-import to the mitochondrial lumen and is essential for cell survival^6,7^.

Several structures determined by X-ray crystallography are available for *Thermotoga maritima* CorA (TmCorA)^8–12^. All wild-type proteins have been crystalized as nearly symmetric pentamers in the presence of divalent metal ions and all represent a non-conducting state of the channel with a narrow and hydrophobic pore. Figure 1A shows a representative structure, which is characterized by a transmembrane domain (TMD) connected to the intracellular domain (ICD) by a long stalk helix. The periplasmic entrance to the pore contains the conserved GMN motif that presumably binds to Mg^2+^ via its first hydration shell and thereby acts as a selectivity filter^11,13,14^. The ICD contains ten inter-protomer binding sites for Mg^2+^ (two per protomer, denoted M1 and M2) involved in regulating the channel^11,15,16^. The open state(s) of CorA have so far not been crystallized, but several biochemical and structural studies^15,16^ as well as molecular dynamics simulations^12^ have pinpointed the determining residues involved in gating and suggested open models. One model suggests pore dilation upon loss of Mg^2+^ at the M1 (and M2) sites due to a concerted iris-like movement^16,17^, while another suggests a hydrophobic-to-polar transition of the pore upon concerted rotation of the stalk helices^18,19^.

**Figure 1.**
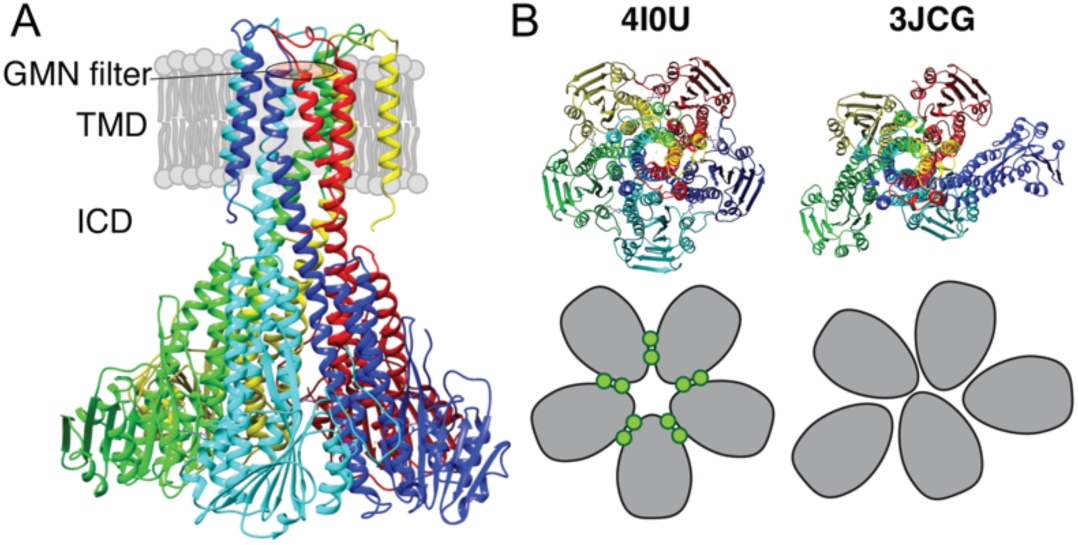
X-ray and cryo-EM structures of CorA. A: Side view of symmetric CorA (PDB ID: 4I0U) in presence of Mg^2+^ (“closed form”). B: Top view of the same symmetric state of CorA (PDB ID: 4I0U) side-by-side to one of the asymmetric states observed in the absence of Mg^2+^ (“open form”) (PDB ID: 3JCG). A schematic representation of the two forms is shown below their structures, with each monomer shown in gray and Mg^2+^ ions represented as green circles.

Recently, cryo-EM structures were obtained both in the presence and absence of Mg^2+^ ^20^. The Mg^2+^ bound structure at ∼3.8 Å resolution was symmetric and closed, in line with crystal structures, whereas two Mg^2+^-free structures at ∼7.1 Å were symmetry broken and with dilated pores. Figure 1B shows an intracellular view of the symmetric and asymmetric states, highlighting the symmetry break upon removing Mg^2+^. From these observations, the proposed model involves a sequential destabilisation of CorA upon Mg^2+^ removal, leading to a highly dynamic protein with shuffling protomers in the ICD, increasing the likelihood of pore dilation and wetting events^20,21^. Recent coarse grained MD simulations revealed the residue level details of how a complex interaction network involving asymmetric movements of ICD monomers ultimately leads to a conducting state upon removal of Mg^2+^ ^22^. High-speed atomic force microscopy (HS-AFM) data on densely packed CorA in lipid bilayers supported this model, but at the same time provided more insight to the dynamic interconversion of different states, including a fourth population of highly asymmetric CorA, not resolved by cryo-EM^23^. Interestingly, this population accounted for most observed conformations at low Mg^2+^ concentrations, supporting that CorA is a dynamic protein with a relatively flat energy landscape and, potentially, multiple open states. However, CorA mutants with mutated regulatory M1 sites were still able to crystallize in the (symmetric) closed state^19^, suggesting that inter-protomer binding of Mg^2+^ is not required for closing the channel. Overall, the cryo-EM and AFM experiments hint towards a highly dynamic ensemble of primarily asymmetric states at low Mg^2+^ concentrations, while the successful crystallisation of M1 site mutants suggests that the closed state is significantly present at these conditions.

In this study, we investigated CorA using two room-temperature methods, namely small-angle neutron scattering (SANS), sensitive to large amplitude conformational changes and magic-angle spinning solid-state NMR (MAS NMR), sensitive to structure and dynamics with atomic resolution^24,25^. For both methods, we employ custom developed state-of-the-art methodology, i.e. size-exclusion chromatography (SEC) coupled to SANS^26,27^ and match-out deuterated carrier systems for SANS^28,29^ (so-called stealth carrier systems), and >100 kHz MAS NMR in lipid bilayers^30,31^. Based on these data in conjunction with molecular simulations and modelling, we propose a model in which CorA is in a dynamic equilibrium between symmetric and asymmetric states, independent of bound Mg^2+^, but where an ensemble of conducting states is energetically more favourable for Mg^2+^-free CorA due to increased conformational dynamics resulting from the released electrostatic constraint.

## Results

### CorA is structurally similar in presence and absence of Mg^2+^

The published cryo-EM structures of CorA in absence of Mg^2+^ (Figure 1B, 3JCG) reveal large structural rearrangements compared to the nearly symmetric, non-conductive state obtained from crystallography (Figure 1B, 4I0U). SANS curves calculated from these two structural states of CorA reveal a significant change in the scattering curve in the region *q* = 0.08 Å^-1^ –0.15 Å^-1^ (Figure 2A and Figure 2B, right panels), i.e. on a length scale that is well-covered in a standard SANS experiment. To match the cryo-EM conditions, we performed SANS measurements in n-dodecyl-B-D-maltoside (DDM) detergent micelles and 2-Oleoyl-1-palmitoyl-sn-glycero-3-phosphocholine (POPC) lipid nanodiscs. We used selectively deuterated versions of both carrier types that were homogenously matched-out and hence invisible at 100 % D_2_O; i.e. stealth DDM (sDDM) and stealth nanodiscs (sND, Figure 2-figure supplement 1). Strikingly, the measured SANS curves are pair-wise indistinguishable in the absence of Mg^2+^ (1 mM EDTA) and in the presence of 40 mM Mg^2+^ for the sDDM (Figure 2A) and sND (Figure 2B) samples, respectively, indicating no significant difference in the average conformations of the Mg^2+^-free and bound states of CorA. This observation contrasts with the recently proposed large-scale structural rearrangements reported from cryo-EM and high-speed AFM data.

**Figure 2.**
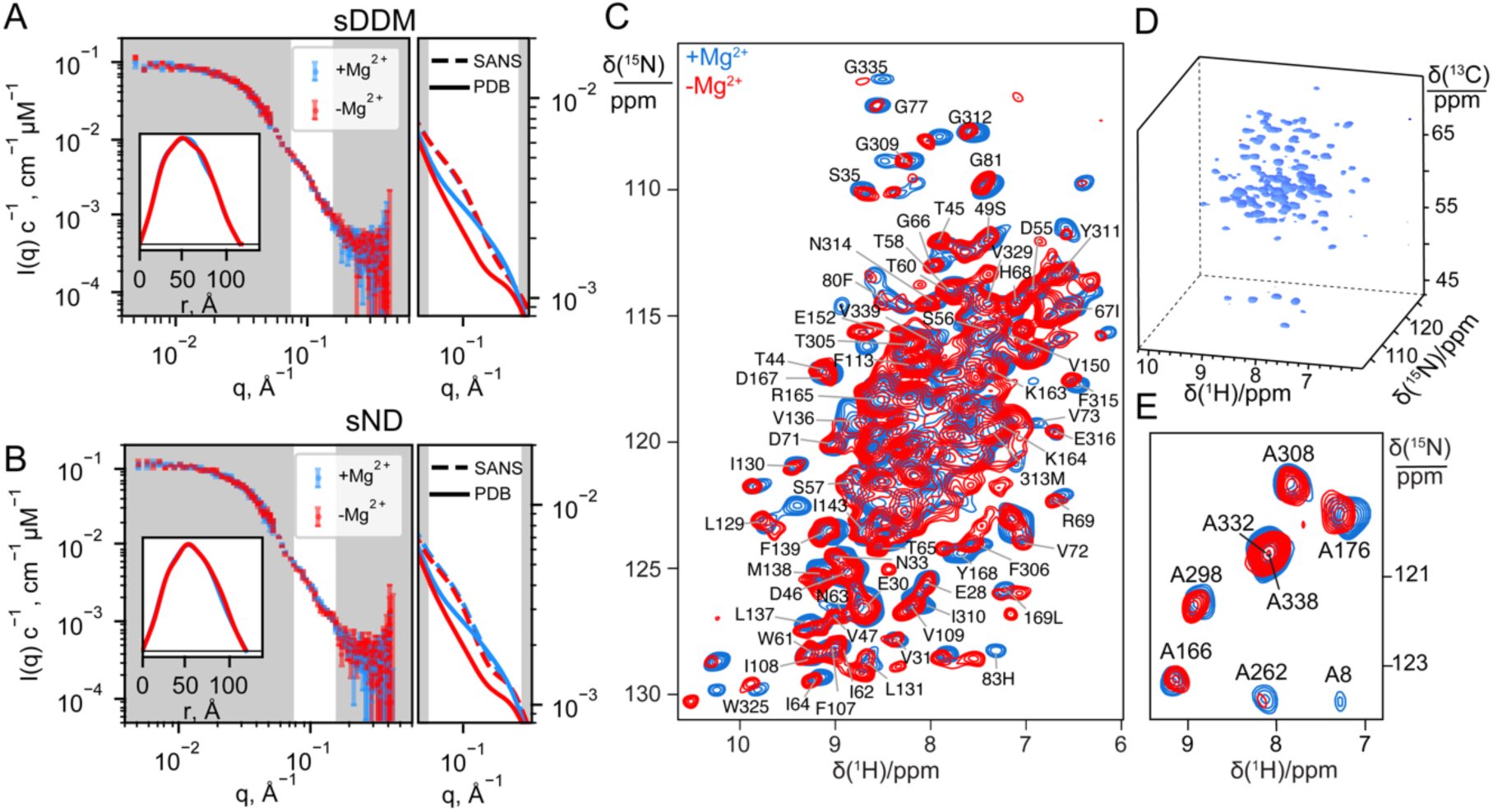
Experimental data on CorA in presence (blue) and absence (red) of Mg^2+^. A+B: Experimental SANS data of CorA embedded in stealth DDM micelles (sDDM) and stealth nanodiscs (sND), respectively, with p(r)-distributions calculated on BayesApp^32^ in the inset. The rightmost plots show zoomed comparisons of the p(r)-fits of experimental data (SANS, dashed lines) with the SANS curves calculated on the X-ray (4I0U) and cryo-EM (3JCG) structures (PDB, full lines). Complete fits based on the PDB structures are shown in Figure 4A. C: 2D ^1^H-^15^N dipolar correlation spectra by MAS NMR of CorA embedded in hydrated DMPC bilayers recorded at 1 GHz ^1^H Larmor frequency and 107 kHz MAS. D: Cube representation of a 3D ^1^H-^15^N-^13^Cα spectrum. E: 2D ^1^H-^15^N dipolar correlation spectra obtained for ^15^N-Alanine labeled CorA recorded at 1 GHz ^1^H Larmor frequency and 60 kHz MAS. In C and E, site-specific assignments are annotated for resolved resonances.

We note that the SANS data obtained on the sND samples (Figure 2B) have a slight excess scattering contribution at low-*q* compared to the sDDM samples (Figure 2A), which can be attributed to the presence of a few *E. coli* endogenous lipids in the sND samples. However, the SANS data from sDDM and sND samples are indistinguishable in the *q*-region expected to reveal differences from symmetric and asymmetric states (Figure 2-figure supplement 2), which confirms that CorA exhibits the same behaviour in a POPC lipid environment and in DDM detergent carriers. The rightmost panels of Figure 2A and Figure 2B show enhanced views on this region for the SANS data compared to the SANS curves calculated from the PDB structures. Interestingly, neither of the curves calculated from the PBD structures match the measured SANS data, suggesting that the solution structure of CorA cannot be described by any of these single structures, regardless of whether or not CorA is in the presence of Mg^2+^.

While SANS data provided information on the overall molecular shape of CorA in the two preparations, we used MAS NMR to obtain insight into structural changes at the residue-level length-scale. MAS NMR data were recorded on uniformly ^13^C,^15^N-labelled CorA, reconstituted in 1,2-dimyristoyl-sn-glycero-3-phosphocholine (DMPC) lipid bilayers, in the presence or absence of Mg^2+^. Backbone resonance assignment was obtained at high Mg^2+^ concentration acquiring a set of three-dimensional experiments relying on ^1^H^N^ and ^1^Hα detection with 100 kHz MAS. We were able to annotate ∼100 peaks to residues spread throughout the structure of CorA. Notably, the assignment of CorA with and without Mg^2+^ is clustered in the globular region in the ICD and in the TMD, including the important periplasmic loop, whereas only sparse assignments were established in the long portion (243-289) of the stalk helix connecting the two regions. The determination of random coil chemical shift deviation (CSD) values confirmed that the secondary structure is in good agreement with the one obtained by X-ray crystallography and Cryo-EM.

2D ^1^H-^15^N dipolar correlation spectra represent direct structural “fingerprints” of CorA in the two preparations. Despite the high signal overlap associated to the high molecular weight of CorA, fast MAS rates and the ultra-high magnetic field guarantee high sensitivity and feature numerous signals with a resolution sufficient to track subtle structural changes. Once again, against our expectations, we remarked that the spectra with and without Mg^2+^ showed very little difference, with the positions of the resolved peaks differing less than 0.1 ppm in the two forms and without peak splitting or broadening that would indicate distinct conformations (Figure 2C).

Two parallel strategies were pursued to extend the analysis to the more crowded regions. First, we acquired three-dimensional (3D) experiments which correlate the amide proton and nitrogen with the C*_α_*-carbon within each residue and thus include an additional ^13^C chemical shift dimension (Figure 2D). The resulting 3D spectra confirmed negligible chemical shift variations over more than ∼90 sites across the TMD and ICD. In the Mg^2+^-free sample, however, a notable decrease in signal intensity was observed for most residues, resulting in the complete disappearance of two thirds of the peaks from the TMD (vide infra).

Secondly, we used amino acid-specific isotopic enrichment to select the NMR signals associated to the amide groups of the alanine residues. Each CorA protomer contains eight alanine residues, distributed with four in the TMD and four in the ICD, which were all visible and assigned in the corresponding ^1^H-^15^N dipolar correlation spectra of two preparations with and without magnesium (Figure 2E). Also, in this case, the spectra are superimposable and show no evidence of peak splitting.

In conclusion, and in line with SANS, the NMR data show that the predominant structure of CorA in lipid bilayers is unaltered by the removal of Mg^2+^.

### CorA is active and preserves its tertiary structure in D_2_O

Since Mg^2+^ hydration plays an important role in CorA selectivity, and D_2_O and H_2_O have slightly different physicochemical properties^35^, we speculated whether the identical SANS curves with and without Mg^2+^ were due to CorA losing its activity in the SANS condition, i.e. at 100% D_2_O. To test this, we measured the activity of CorA in POPC liposomes under the SANS conditions by a fluorometric assay (Figure 3A). This analysis shows that CorA is clearly active in D_2_O. The transport rate estimated from the initial linear part of the trace is lower by less than a factor of two compared to H_2_O. A slightly reduced rate in D_2_O has been reported for other membrane proteins^36^ and is explainable by slightly altered properties of the two solvents^35,37^. We could also inhibit CorA activity in D_2_O with Co[NH3]_6_^3+^ (Figure 3A, green), supporting that the protein is indeed functional under the SANS conditions.

**Figure 3.**
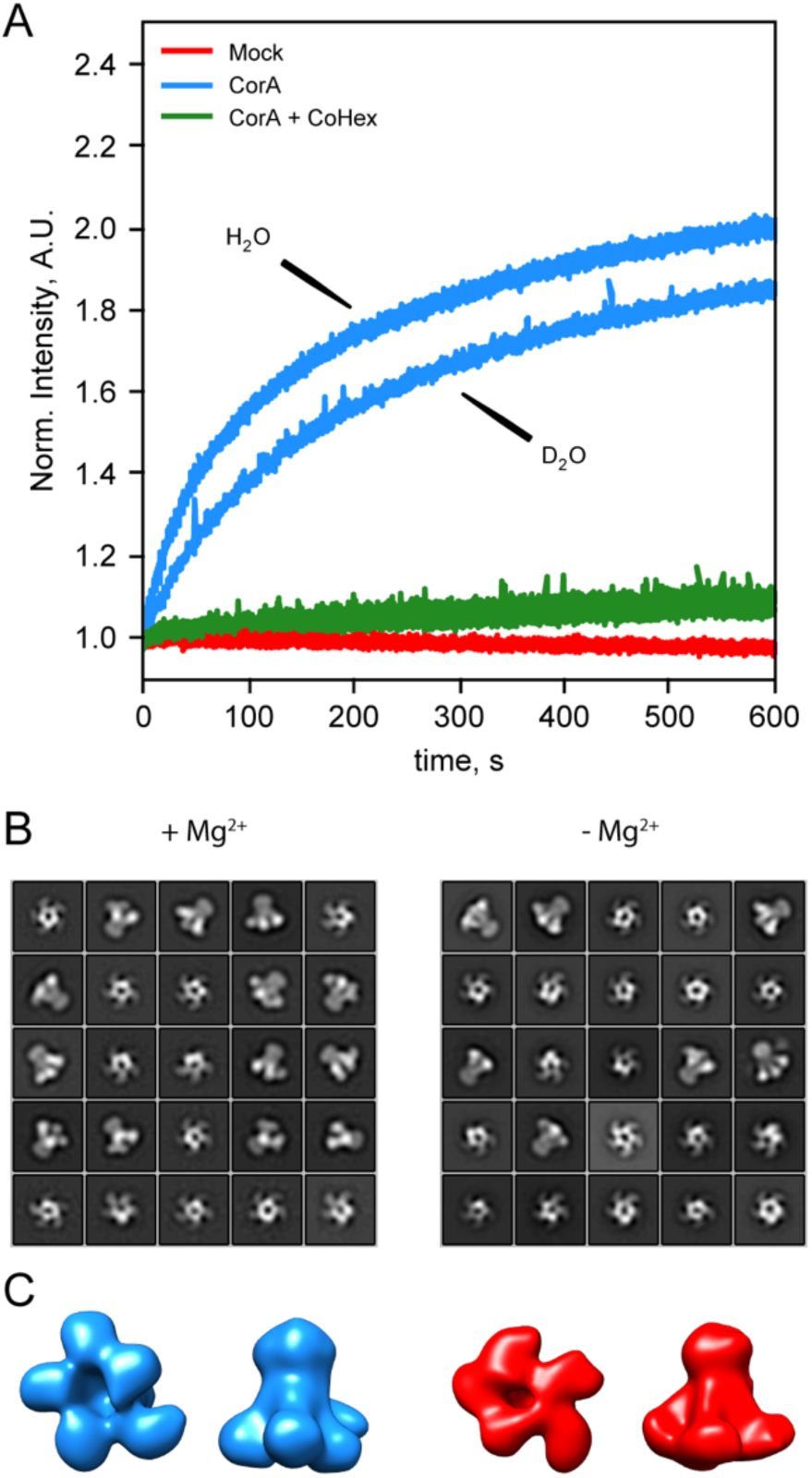
CorA activity in D_2_O and direct visualization by negative stain EM. A: CorA activity in the conditions used for SANS. The traces are the normalized fluorescence signals after adding Mg^2+^ to either empty POPC liposomes (Mock), CorA-POPC proteoliposomes (CorA), or CorA-POPC proteoliposomes preincubated with the inhibitor Co[NH_3_]^3+^ in D_2_O (CorA + CoHex). B: Negative stain EM of CorA in DDM and D_2_O with and without Mg^2+^. The 25 most abundant 2D classes are shown for each condition. The box dimensions are 170 x 170 Å^2^ for scale. C: Final 3D model for each condition shown from the intracellular side and in side-view, respectively.

We further carried out negative stain EM on samples of CorA in DDM and D_2_O with or without Mg^2+^. The refined 2D classes clearly show that the pentameric architecture of CorA is preserved in D_2_O in both conditions (Figure 3B). Furthermore, several 2D classes appear to exhibit approximate fivefold symmetry, both in the presence and absence of Mg^2+^. To avoid bias, we refined 3D models without imposing symmetry (Figure 3C). In both conditions, these low-resolution 3D models (≈ 15 Å) are reminiscent of the overall expected architecture of CorA (Figure 1), but notably show some degree of asymmetry. In conclusion, CorA show little to no perturbation from measurements in D_2_O, rendering the SANS data sets viable for modelling of the solution structure of CorA.

### Model refinement to SANS data shows that CorA is asymmetric

The SANS data sets obtained in sDDM (Figure 2A) exhibit well-defined Guinier-regions and the calculated radii of gyration, R_g_, of 42.1 ± 1.3 Å (+ Mg^2+^) and 43.8 ± 1.7 Å (- Mg^2+^) are close to the predicted values from the X-ray (41.3 Å) and cryo-EM structures (42.2 Å and 42.3 Å). Thus, these data are indicative of well-separated CorA pentamers with no interference from visible lipids or the kind, providing the optimal basis for structural modelling. Given the identicality of the SANS data obtained on the samples of CorA in sDDM with and without Mg^2+^, we performed structural modeling on only a single data set, that is CorA in sDDM without Mg^2+^ (Figure 2A, red). Despite controversies on the open state, there is consensus that the crystallized symmetric state represents the closed state of the protein. Surprisingly, we could not obtain good fits of the symmetric state to our SANS data without clear systematic deviations, especially at the feature present at *q* ≈ 0.1 Å^-1^ (Figure 4A, 4I0U). This was also the case for the asymmetric cryo-EM structure (Figure 4A, 3JCG) that produced an even worse fit. In SANS, the signal represents a population-weighted average of all conformations that the protein can adopt. With a measurement time on the order of several minutes and illumination of ≈ 10^12^-10^13^ molecules, all accessible populations are expected to contribute to the signal. A relatively flat energy landscape with multiple interconverting states has been proposed in Mg^2+^-free conditions, making a fit of a single structure less meaningful in this context. However, it is unlikely that the average of an ensemble of asymmetric structures give rise to the same SANS signal as that of a single symmetric state corresponding to the structure determined by crystallography and cryo-EM in high Mg^2+^.

**Figure 4.**
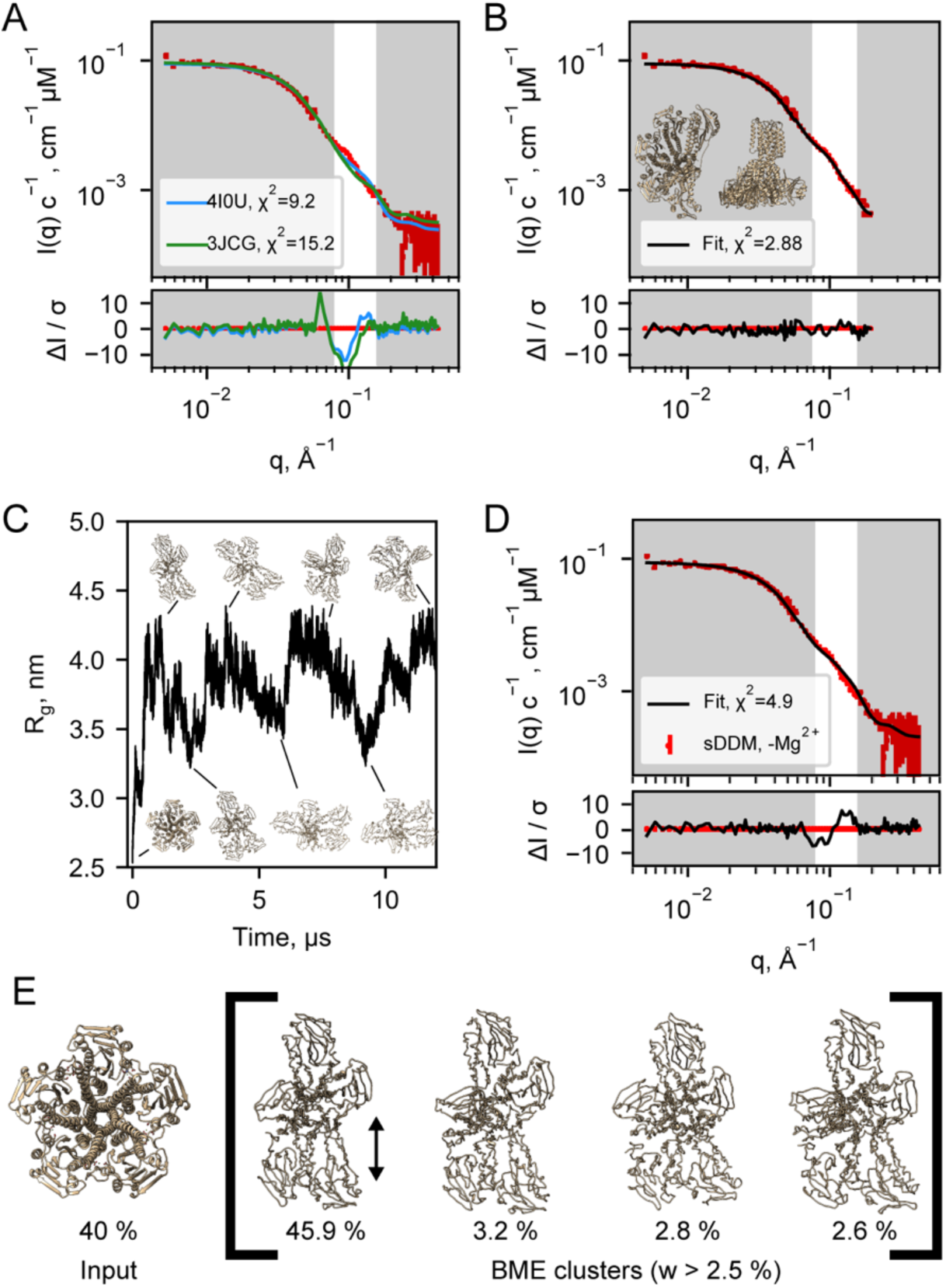
Structural modeling of CorA from SANS data. A: Model fits of the closed crystal structure (4I0U) and open cryo-EM structure (3JCG) to the experimental SANS data obtained in sDDM without Mg^2+^. The bottom panel shows the error-normalised difference plot of the fits. B: Model fit of the structure obtained by regularised normal mode analysis (bottom and side views inserted on plot). C: MetaD molecular dynamics trajectory with representative frames visualized as ribbon structures. D: Fit of a weighted ensemble obtained from the MD simulation in B together with a 40 % contribution from the symmetric CorA. E: The 4 highest weighted cluster centroids in the metaD simulation (right bracket) together with the closed symmetric structure (left) which together illustrates the ensemble of CorA structures that produced the best fit to the experimental SANS data.

With no scattering contribution from the carrier systems, it becomes possible to analyse the SANS data by conventional methods for soluble proteins, such as bead-modeling. When imposing P5 symmetry, we could obtain envelopes reminiscent of the CorA structure by bead-modelling (Figure 4-figure supplement 1B), whereas no symmetry (P1) imposed lead to asymmetric mass distributions that were not at all compatible with the overall architecture of CorA (Figure 4-figure supplement 1A). This indicates that there is significant structural dispersion in the sample.

To obtain a molecular constrained model compatible with the data, we applied a modified type of normal mode analysis (NMA), starting from the closed crystal structure. A structure with mostly intact secondary structure but a high degree of asymmetry in the ICDs yielded a great fit to the SANS data (Figure 4B). Importantly, this model describes the feature at *q* ≈ 0.1 Å^-1^, where the PDB models deviate the most. Thus, on average, the solution structure of CorA appears to be asymmetric, in line with our EM models (Figure 3C). Again, we emphasize that such an overall asymmetric structure of CorA in presence of excess Mg^2+^ is in stark contrast to the picture of a closed, symmetric structure that has served the basis for all proposed mechanisms of Mg^2+^ gating. However, a single asymmetric model as derived from NMA (Figure 4B) is neither compatible with a single set of peaks in NMR (Figure 2C and Figure 2E) nor the substantial experimental evidence for a nearly symmetric, closed state in presence of Mg^2+^ . Likely, CorA adopts multiple different states^20,23^, and according to our data does so both with and without Mg^2+^ bound. In this case, the SANS data would represent the number weighted average of different states that must be overall asymmetric.

To model the apparent asymmetry in CorA in more detail, we performed coarse grained molecular dynamics simulations (MD). First, we set up CorA embedded in a POPC bilayer using the Martini3.0b force field. Starting from the symmetric or asymmetric structures, 32 µs and 20 µs simulations, respectively, without any inter-chain elastic network terms yielded only small structural fluctuations (data not shown), which did not significantly improve fits to the experimental SANS data, especially not around the feature at *q* ≈ 0.1 Å^-1^. Thus, we extended the analysis to metadynamics simulations (MetaD) that allows for enhanced sampling of structural dynamics. MetaD drives the simulation towards a larger variety of structural states based on an energetic bias on a structural feature, a so-called collective variable, here the R_g_ on specific ICD residues. Starting from the symmetric structure, the MetaD simulation quickly drove the simulation away from the local structural minimum that the standard MD had been trapped in and sampled a large range of conformationally different structures (Figure 4C).

The averaged back-calculated SANS of the entire ensemble of structures obtained from MetaD simulations did not fit the experimental SANS data satisfactorily (data not shown). This can be explained by inaccuracies in the simulation that arise from e.g. inaccuracies in the coarse grained force field and imprecisions in the simulation that in turn arise from e.g. insufficient sampling. To resolve this, we applied the Bayesian/Maximum Entropy reweighting method (BME) to optimize the weights of the individual conformations in the simulation with the aim of obtaining an ensemble in better agreement with the experimental SANS data. In addition to the BME, we enforced that a symmetric state should be present in the final ensemble, given the substantial experimental evidence for this state in the literature and that it was under-sampled in the simulation. The best fit to the SANS data (Figure 4D) was obtained with an ensemble consisting of 40% ± 28 % symmetric CorA and the remainder of asymmetric conformations from the MetaD (Figure 4E and Figure 4-figure supplement 3). Despite some systematic deviations, the fit is much improved with regards to describing the feature in the SANS data at *q* ≈ 0.1 Å^-1^, as compared to the fits obtained with the symmetric crystal structure or the open cryo-EM structures, respectively (Figure 4A).

To visualise the results of the MetaD simulation and hence the simulated dynamics, we cluster similar structures and show the four most predominant cluster centroids (Figure 4E) where especially one cluster predominate in the final fitted ensemble. Although the four cluster centroids are wide apart in simulation time (≈500 ns), they are structurally similar with a maximum pairwise RMSD of 6.5 Å (data not shown). The main difference is the distance between the two protomers (Figure 4E, black arrow), which indicate that the individual domains of the ICD can move relative to each other. This is in line with the subunit displacements described in the symmetry-break based gating model derived from the cryo-EM structures and supported by AFM measurements. Importantly, however, we find that these movements occur irrespective of the presence of Mg^2+^. Interestingly, CorA mutants were recently shown to crystallize in the symmetric, closed state without Mg^2+^ bound at the regulatory M1 sites. While the channel is surely closed in the presence of high Mg^2+^, this observation challenges the claim that release of interfacial bound Mg^2+^ is the cause for symmetry-break and opening of the channel at low Mg^2+^ levels. In light of this, our data suggest that CorA could also be in an asymmetric, closed state at high Mg^2+^ levels. Indeed, we find that the structure of CorA is better described as a distribution of symmetric and asymmetric structures with no large-scale conformational differences between the open and closed states.

### Structural dynamics are different in open and closed states of CorA

So far, we have considered a set of static snapshots to interpret the wide variety of populated states of CorA. The observation of MAS NMR dynamical probes sheds light on the backbone motions of these states over different timescales, enriching the structural description of CorA with conformational plasticity.

A first insight on site-specific dynamics is obtained by comparing the peak intensities observed in the MAS NMR experiments in the two samples with and without Mg^2+^.

Peak intensities are dependent on dipolar couplings between nearby nuclei and are affected when such couplings are averaged by local motions. For an amide ^1^H-^15^N pair, this corresponds to motional processes faster than ∼ tens of kHz (i.e. more rapid than ∼ hundreds of μs).

Changes in peak intensity are already noticeable in the 2D ^1^H-^15^N dipolar correlation spectra, but are amplified in the 3D correlations, where additional radio-frequency irradiation periods act as a more stringent filter, dumping the signals of the most mobile sites. 1D traces of two exemplar 3D ^1^H-^15^N-^13^C correlations (T65 in the ICD and F306 in the TMD) and the plot of signal intensities over the full protein sequence with and without Mg^2+^ are shown in Figure 5A and Figure 5B, respectively. As mentioned above, an overall decrease in peak intensities is associated to removal of Mg^2+^, with a stronger effect observed in the TMD. This points toward a variation of the dynamic behavior of this region in the two samples.

**Figure 5.**
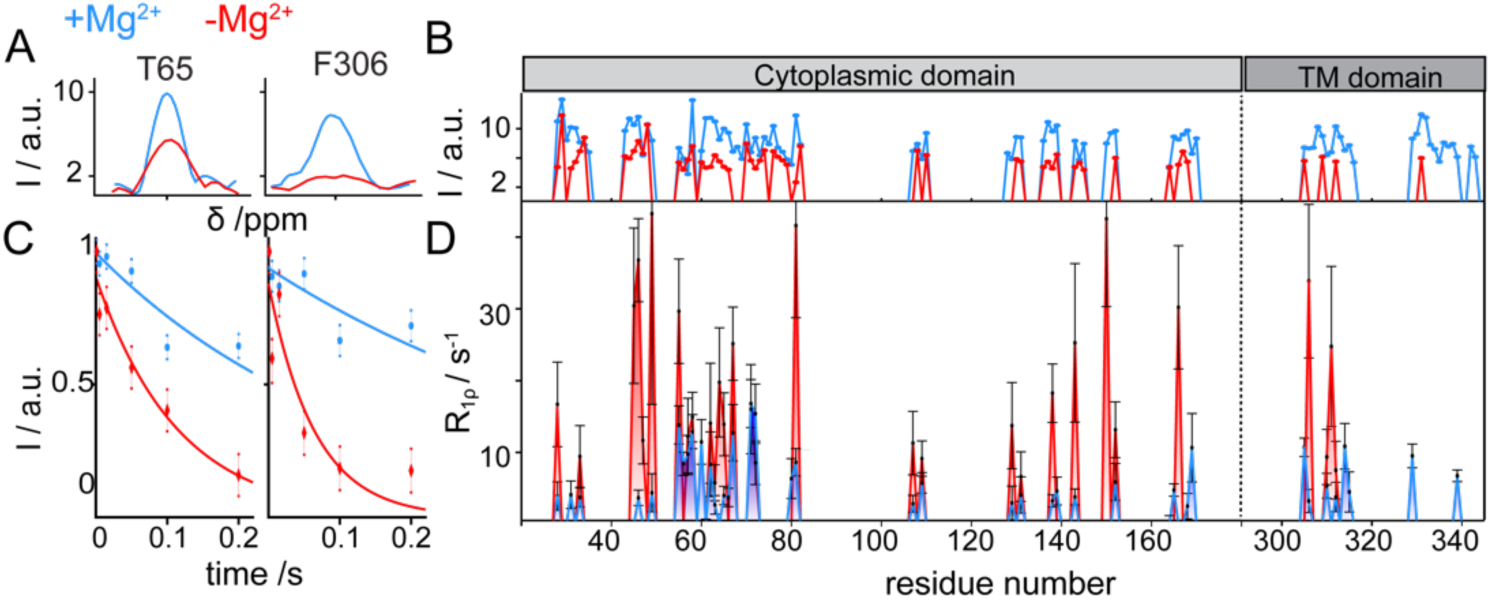
CorA backbone dynamics by MAS NMR in DMPC in presence (blue) and in absence (red) of Mg^2+.^ A: Examples of 1D traces of 3D ^1^H-^15^N-^13^C peaks for two residues in the ICD (T65) and in the TMD (F306). B: Comparison of peak intensities in 3D ^1^H-^15^N-^13^C spectra over the protein sequence. ICD and TMD are indicated by boxes of different color on top of the plot. C-D: Site-specific ^15^N R_1ρ_ rates measured with a 15 kHz spin-lock field. C: Examples of ^15^N R_1ρ_ relaxation decays together with the corresponding mono-exponential fits for residues T65 and F306. D: Comparison of site-specific backbone ^15^N R_1ρ_ rates plotted along the CorA sequence.

^15^N spin-lattice relaxation rates in the presence of a spin-lock field (^15^N R_1ρ_) are sensitive reporters of motions occurring over a window of hundreds of ns to hundreds of μs. These parameters were measured site-specifically for ∼40 residues in CorA backbone, by introducing a relaxation filter in the ^1^H-^15^N dipolar correlation module and monitoring the signal decay of each amide pairs in a series of experiments with increasing relaxation delays (Figure 5C). A remarkable (up to 10 fold) increase was observed throughout the entire protein upon removal of Mg^2+^ (Figure 5D). The same effect was observed for the ^15^N-alanine labelled sample (Figure 5-figure supplement 1A). An additional measurement of ^15^N R_1ρ_ rates upon a ∼20°C cooling revealed a different temperature-dependent behaviour of the two Mg^2+^-loaded samples (Figure 5-figure supplement 1B). While in the presence of Mg^2+^ relaxation rates were conserved at low temperatures, an important (2-3 fold) decrease was observed in the Mg^2+^-free form. Finally, also dynamics on the ns timescale obtained by the measurements of bulk backbone ^15^N R_1_ showed a global increase of relaxation rates (by a factor ∼1.5) upon removal of Mg^2+^ (Figure 5-figure supplement 1C).

In summary, MAS NMR reveals that removal of Mg^2+^ triggers an increase in the backbone flexibility on different timescales, a dynamical effect which is different for the ICD and the TMD.

## Discussion

The functional mechanism of the pentameric divalent cation channel CorA of *Thermotoga maritima* has been under investigation since the release of the first high resolution structures in 2006. It is now well acknowledged that gating involves a conformational transition between a closed symmetric state and one or many asymmetric states. The concise current model derived from cryo-EM^20^ and HS-AFM^23^ defines a rigid symmetric conformation at high Mg^2+^ concentrations (>4 mM), dynamic asymmetric conformations at low Mg^2+^ concentrations (2-3 mM) and several distinct rigid asymmetric conformations in the absence of Mg^2+^.

We have performed SANS in invisible carrier systems to study the solution conformation of CorA, together with modelling of both static and ensemble structures obtained from MetaD simulations. Furthermore, we used ^1^H-detected solid-state NMR to investigate CorA conformation and dynamics in hydrated lipid bilayers. These complementary methods allow us to expand the current view on the mechanistically important conformational equilibria.

### Asymmetric states are populated in CorA at high Mg^2+^ concentrations

Crystal structures of CorA from *Thermotoga maritima* (*Tm*) and *Methanocaldococcus jannaschii* (*Mj*) solved in the Mg^2+^-bound state revealed a symmetric bell-like structure, showing a narrow inner channel, representing the closed form. All proposed gating models were built on the common basis of a single rigid and symmetric conformation at high Mg^2+^ concentrations^11,12,16,20,23,40,41^. Our SANS data recorded on CorA in high Mg^2+^ concentrations is inconsistent with a single symmetric structure and can only be explained by symmetry breaking and/or a mixture of populations e.g. through the concomitant presence of symmetric and asymmetric states. Indirect indications for a mixture of populations even in the presence of Mg^2+^ can be found in the literature: For example, the symmetric cryo-EM structure (3JCF) obtained at high Mg^2+^ concentrations, resulted from roughly 60% of the particles^20^, which implies that asymmetric states could partly account for the remaining 40%. Although with lower resolution, our negative-stain TEM data indeed support the presence of such populations.

HS-AFM measurements^23^ showed that the symmetric, closed state was stable in starting conditions at high Mg^2+^. However, only 35 % of this state was recovered after complete removal and reintroduction of Mg^2+^ ions, suggesting secondary effects working on adjusting the equilibrium between fluctuating asymmetric states and the closed symmetric state.

### A symmetric state is populated in CorA in absence of Mg^2+^

The conformation of metal-free CorA has been the object of a long ongoing speculation since wild-type CorA failed to crystallize in the absence of Mg^2+^. Asymmetric subunit arrangements were first proposed based on an X-ray structure of a truncated CorA variant (ΔN25/R222A/K223A)^12^. Since the release of two cryo-EM structures of wild-type CorA^20^, it is now a common belief that this channel adopts exclusively an asymmetric arrangement in absence of Mg**^2+^**.

NMR spectra are sensitive to the environments of each nucleus, and thus report on conformational transitions breaking the local symmetries. The fact that the NMR fingerprint spectra do not change significantly upon removal of Mg**^2+^** indicates the conservation of local symmetric environments for most of the NMR-active nuclei in both conditions. This evidence in turn suggests that the global symmetry is maintained for a substantial population of CorA pentamers, as it is unlikely that the ample rearrangements implied by the cryo-EM model could occur without affecting local symmetries of so many nuclei in the five subunits. This view is additionally corroborated by our negative-stain TEM data, which showed the presence of nearly symmetric 2D classes in Mg**^2+^**-free preparations.

Symmetry breaking is expected to result in local conformational heterogeneities between the monomers, resulting in peak splitting, with a possible reduction of the intensity beyond the detection limit. In this regard, the general decrease in intensity of the NMR spectra observed in absence of Mg**^2+^** suggests a significant reduction of the population of the observed symmetric state, compatible with the co-existence of a broad distribution of asymmetric states. It has previously been shown that CorA mutants with abolished M1 regulatory sites still crystallize in a symmetric, closed state. This suggests that binding of Mg^2+^ is not necessary for closing the channel^19^.

The two asymmetric cryo-EM structures were refined from only about 15 % each of the picked particles, indicating the presence of various conformations, and possibly the symmetric form. AFM provided a more detailed conformational analysis and found around 20% symmetric structures at low Mg^2+^ concentrations (0-3 mM). The closely related Zn transporter ZntB also showed a symmetric structure in the absence of regulatory cations^42^. All these observations point towards a conformational equilibrium between the symmetric and various asymmetric states which is partially but not entirely shifted towards asymmetric forms at low cation concentration.

### Increased dynamics in CorA in absence of Mg^2+^

A “dynamic character” of CorA at low levels of Mg^2+^ was postulated on the bases of the cryo-EM data^20^ and recently characterized by high-speed AFM^23^. An elevated conformational plasticity would indeed be consistent with the hitherto unsuccessful crystallization of a conducting state.

MAS NMR has the exclusive advantage of probing molecular motions with site-specific resolution. Differently from a previous NMR study on a truncated construct in detergent micelles^43^, we here directly tackled the full-length protein in lipid bilayers. Notably, we characterized the change in dynamics occurring upon Mg^2+^ release by acquiring NMR observables sensitive to different timescales. On a fast ps-ns timescale, increased bulk ^15^N R_1_s in the absence of Mg^2+^ are indicative of less restricted backbone motions, as also suggested by MD simulations^41^. Site-specific ^15^N R_1ρ_ rates are reporters of slower segmental motions displacing secondary structure elements with respect to each other, over a window of hundreds of ns to hundreds of μs. These dynamical processes appear to be largely promoted over the whole CorA structure when Mg^2+^ is absent.

Notably, a change in dynamics is also visible in the TMD region. Here we detected and assigned most trans-membrane and periplasmic residues including the GMN motif in the presence of Mg^2+^. However, for these residues, peaks broadened beyond detection when Mg^2+^ was removed. This points towards the presence of multiple conformations in this region as well, although the exchange dynamics likely occurs on a different regime with respect to the ICD domain. An earlier work suggested that removal of Mg^2+^ results in a combination of lateral and radial tilting of two adjacent monomers, which allows the creation of interactions between them^12^. This idea was recently extended based on coarse-grained MD simulations that show multi-step conformational changes propagating from the ICD to the TMD helices^22^. In line with these observations, we propose that collective ns-μs motions of the backbone initially promoted in the ICD by release of Mg^2+^ ions would in turn induce higher conformational flexibility in the TMD.

### Are all asymmetric states open/conductive? What is an open/conductive state?

Since asymmetric structures are present both in presence and absence of Mg^2+^, lack of symmetry is not equivalent to conduction. The regulation of Mg^2+^ transport clearly needs to be dependent on more stringent control. Interestingly, a previous MD study^21^ found that dry and intermediate transiently wetted states, both non-conducting, were present both with and without bound Mg^2+^ and interconversion rates on the ns time scale. Furthermore, in absence of Mg^2+^, a conducting “stably superhydrated” state was sampled, i.e. where the region of pore lining residues M291, L294, A298, and M302 that form the main hydrophobic constriction is wetted by more than 20 water molecules. Though this state was only sampled in a minor part of the simulations, its population could change at a longer time scale. Indeed, the ns sampled by the simulations cannot describe our observations that CorA adopts symmetric and asymmetric conformations, both in presence and absence of Mg^2+^, while in absence of Mg^2+^, the channel has higher motional freedom on the µs timescale. In light of the MD results, our combined findings support the idea that the “stably superhydrated” state is present in low abundance and interconverts rapidly with the remaining ensemble of states.

### An integrated view on CorA transport regulation

Our findings cannot be explained in terms of sharp transitions between open and closed states. An alternative model that integrates our findings with the previously available literature is schematically illustrated in Figure 6. CorA samples symmetric and asymmetric conformational states, whose distribution is tuned by the Mg^2+^ concentration. Between the symmetric and open states, a relatively flat energy landscape exists with asymmetric, closed states, both in the presence and absence of Mg^2+^. Low Mg^2+^ intracellular levels induce a reduction in the population of the symmetric, closed state together with a decrease of the energy barrier towards the open state, which becomes more populated. At these Mg^2+^ concentrations, CorA can visit a conducting asymmetric ensemble of states. In this context, increased dynamics, as observed by NMR, can become the key determinant allowing CorA to explore different wells of the energy profile, making the open state reachable. Such a dynamic model is equally compatible with previously unexplainable symmetric crystal structures of M1-binding site mutants in the absence of Mg^2+^ ^19^. While for WT CorA, removal of Mg^2+^ increases the dynamics and shifts the conformational equilibria, point mutations probably have an opposite effect, stabilizing the symmetric state and allowing crystal formation.

**Figure 6.**
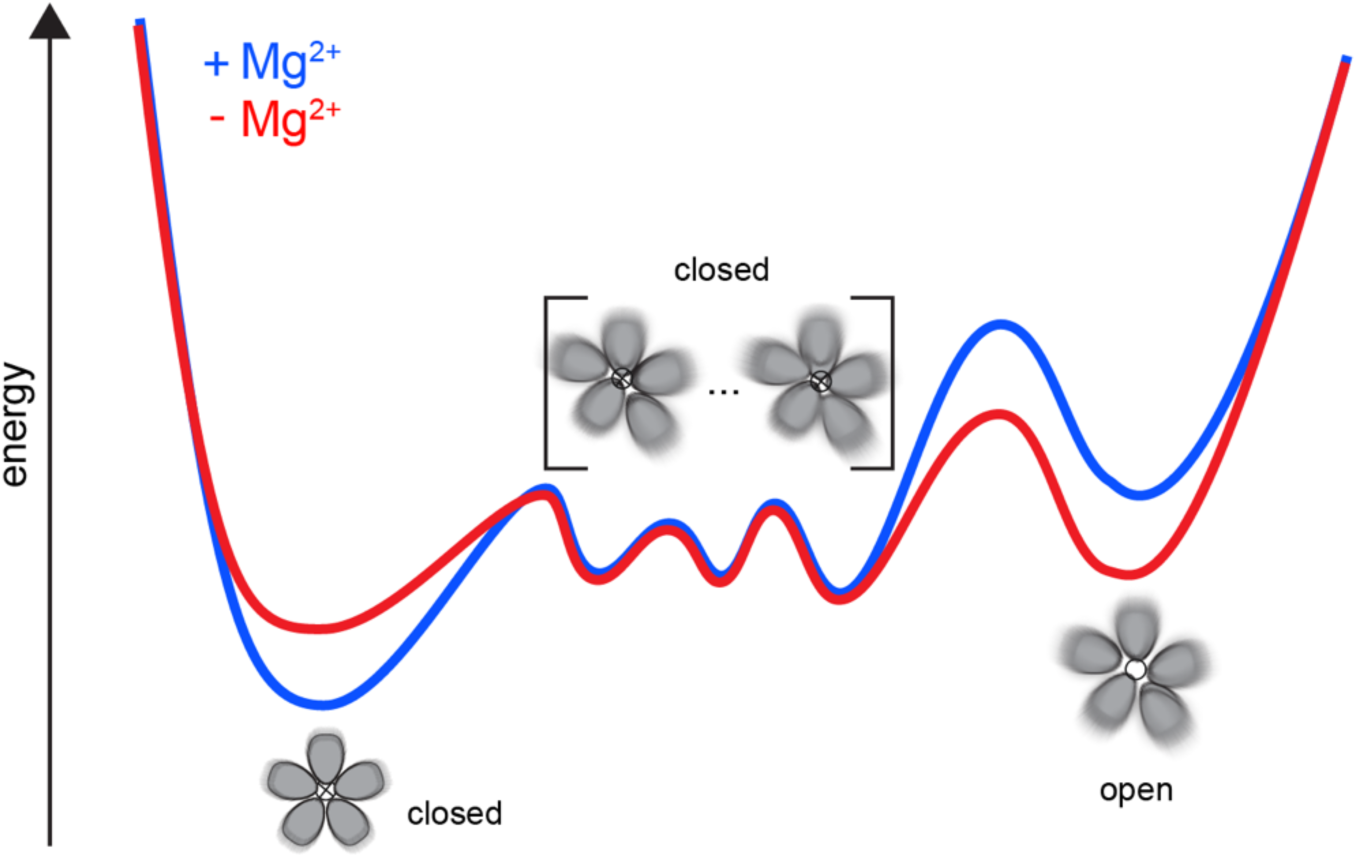
Schematic representations of a dynamic model for CorA. A complex landscape connects a symmetric and closed state with an asymmetric open state, both in the presence and in the absence of regulatory ions.

### Relating to other classes of proteins and channels

Gating has been investigated extensively in the family of pentameric ligand-gated ion channels (pLGIC)^44^, which in general have symmetric or pseudo-symmetric structures in the ligand-free form. Intriguingly, the existence of asymmetric conformations has been observed in the absence of ligand for the serotonine receptor 5HT3R, revealing conformational plasticity also in the resting state. Such conformations are more energetically favorable when the agonist is bound, suggesting an asymmetric activation where resting asymmetric states are possible intermediates of the gating^45^. Asymmetric activation was proposed also for the nicotinic receptor nAChR, where the movement of pore lining helices was shown to induce the asymmetry in the ligand-bound form^46^. Taken together, these studies suggest that asymmetry might have a more widespread occurrence in channel regulation. Asymmetric resting states may exist as intermediates in the conformational landscape independently of the presence of substrate, but they become more energetically favorable when this is bound.

## Conclusion

The availability of room temperature SANS and NMR data of CorA in lipid bilayers, reporting on both global and local behaviour of this channel, supported by MD simulations, allowed us to extend and rationalise the “symmetry-break-upon-gating” model for Mg^2+^ transport. Our observations support the suggestion that asymmetric conformations are involved in the gating mechanism, but in a more complex way than a simpler two-state picture, where Mg^2+^ bound CorA is a stable, symmetric structure. Indeed, we find that the CorA pentamer is a symmetry-broken fluctuating structure able to explore a wide conformational landscape both in the presence and absence of Mg^2+^. We propose that the determining factor for CorA to visit conducting states is the increase in backbone flexibility on different timescales upon release of regulatory Mg^2+^. Future investigations of the conformational equilibria will enrich the insight provided in this study with a better mechanistic understanding of the kinetics and thermodynamics of gating and transport.

## Materials and methods

### Materials

All chemicals were from Sigma-Aldrich unless otherwise stated. DDM was from Carbosynth (UK), match-out-deuterated DDM (sDDM^29^) and match-out deuterated POPC (d-POPC) were synthesised at the National Deuteration Facility at ANSTO (Lucas Heights, Australia). The d-POPC was synthesised as previously^47^, but with custom deuteration (94% D in tail groups, 71 %D in head group). Details of the synthesis and chemical and isotopic analysis is described below.

### Protein production and purification

For SANS, CorA was produced and purified essentially as described elsewhere^48^. For studies in DDM, the N-terminal His-tag was cleaved by tobacco etch virus (TEV) protease before gel filtration, whereas for nanodiscs, it was cleaved after incorporation (see below). For NMR, uniformly isotopically labelled CorA was produced in M9 medium containing 3 g/L ^13^C-glucose and 1 g/L ^15^N-ammonium chloride. ^15^N-alanine labelled CorA was produced in M9 medium containing regular ^12^C-glucose and ^14^N-ammonium chloride and supplemented with 200 mg/L of ^15^N-alanine. Match-out deuterated circularised membrane scaffold protein (MSP), d-csMSP1E3D1, was produced at the D-lab at ILL (Grenoble, France) and purified as described previously^48^. Proteins were stored at -80 °C until used.

### Synthesis of match-out deuterated POPC

The overall synthesis of POPC-*d*_77_ is reported elsewhere^47^. Figure 7 shows the synthetic scheme followed in this study to produce the specific level of deuteration in the head and tail groups of the POPC. The specific level of deuteration in the tail group was achieved by diluting pure heavy water with light water in specific ratios in the Parr reactor when making the deuterated alkyl chains from their fatty acid precursors^29^. The analysis data and spectra of the intermediate compounds and the final compound are shown in Figure 7-figure supplements 1 to 23. Electrospray ionisation mass spectra (ESI-MS) were recorded on a 4000 QTrap AB SCIEX Mass Spectrometer. The overall percent deuteration of the molecules was calculated by ER-MS (enhanced resolution – MS) using the isotope distribution analysis of the different isotopologues by analysing the area under each MS peak which corresponds to a defined number of deuterium atoms. The contribution of the carbon-13 (natural abundance) to the value of the area under each [X+1] MS signal is subtracted based on the relative amount found in the protonated version. In a typical analysis we measure the C-13 natural abundance contribution by running ER-MS of the protonated version (or estimate it by ChemDraw software) and use this value in our calculation using an in-house developed method that subtracts this contribution from each MS signal constituting the isotope distribution. ^1^H NMR (400 MHz), ^13^C NMR (100 MHz), ^31^P NMR (162 MHz) and ^2^H NMR (61.4 MHz) spectra were recorded on a Bruker 400 MHz spectrometer at 298 K. Chemical shifts, in ppm, were referenced to the residual signal of the corresponding solvent. Deuterium NMR spectroscopy was performed using the probe’s lock channel for direct observation.

**Figure 7.**
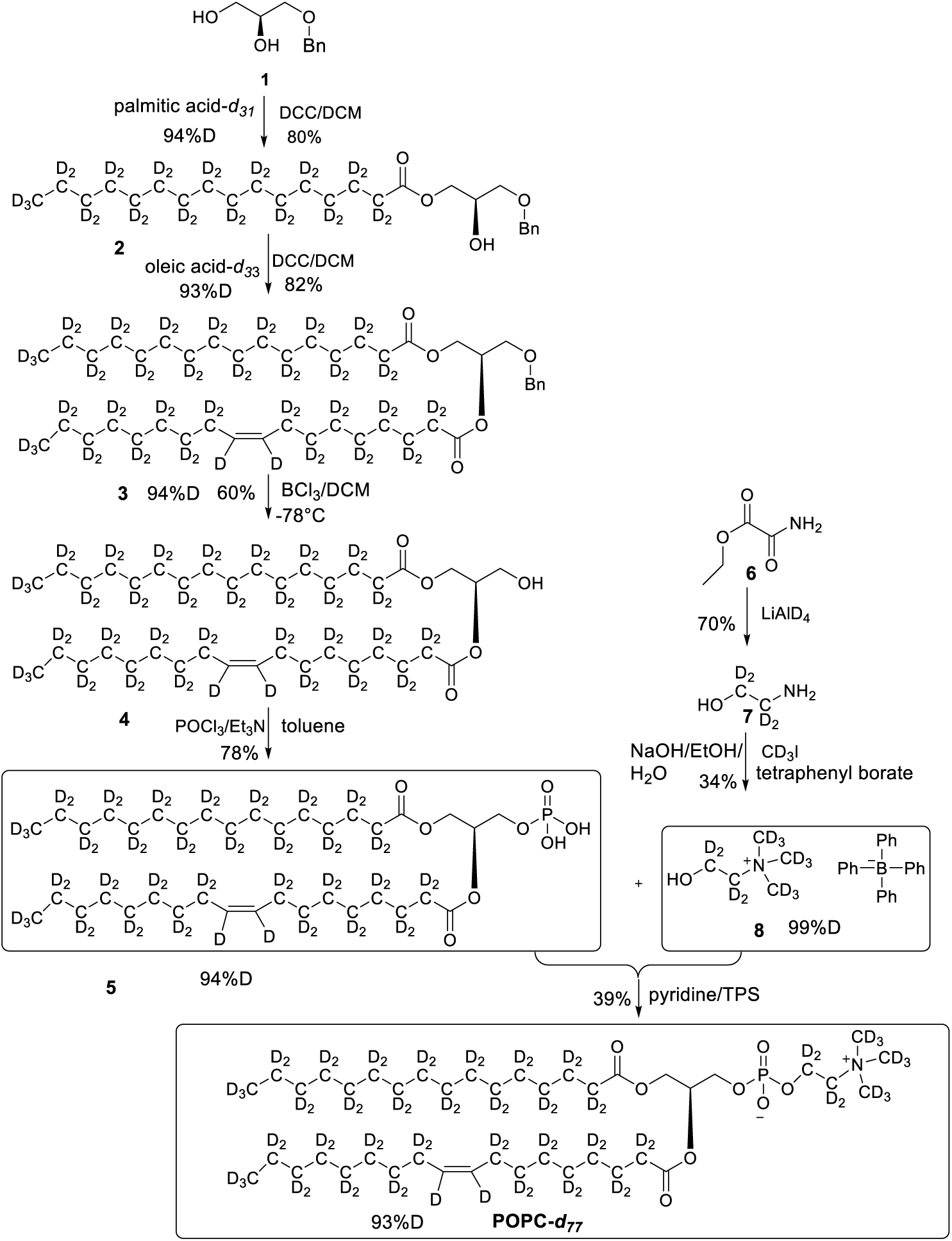
Overall synthesis achieved by following a reported paper, all the intermediates and final POPC-d_77_ were obtained in similar yields^47^.

### CorA reconstitution in sND for SANS

CorA, d-csMSP1E3D1, and d-POPC solubilized in cholate were mixed in a ratio of 400:4:1 with a final d-POPC concentration of 10 mM. Cholate and DDM were removed by adding 50 % w/v amberlite XAD-2 overnight at 5 °C. CorA-loaded sNDs were purified by IMAC on NiNTA resin, added TEV protease and dialysed for three hours at RT followed by ON dialysis at 4 °C against 20 mM TrisHCl pH 8, 100 mM NaCl, 0.5 mM EDTA, 1 mM DTT. Finally, the cleaved sample was purified by reverse IMAC on NiNTA resin. The sample was concentrated to approximately 30 µM before the SANS experiment.

### Reconstitution into multilamellar vesicles for NMR

CorA was reconstituted into DMPC at a lipid-to-protein ratio of 0.5 (w/w) by dialysis against 10 mM Tris-HCl pH 8, 40 mM MgCl_2_, 1 mM methyl-β-cyclodextrin using a 25 kDa MWCO membrane at RT. A white precipitate of multilamellar vesicles was visible after 48 hours and collected by centrifugation after 72 hours. Samples were packed into 1.3 mm or 0.7 mm MAS NMR rotors using an ultracentrifuge tool (Giotto Biotech) at 100.000*g* for 1 hour.

### SEC-SANS

Samples were measured at the D22 beamline at ILL, using the SEC-SANS mode described elsewhere^26,27^, but with the upgrade that UV absorption was measured on the flow cell in the same place, but perpendicular to the neutron beam. The setup was placed in a temperature-controlled cabinet at 11 °C. CorA in sDDM was run on a Superdex200 Increase 10/300 GL column (GE Healthcare) in 20 mM TrisDCl pH 7.5, 150 mM NaCl, 1 mM DTT, 0.5 mM sDDM in D_2_O with an initial flow rate of 0.3 ml/min to allow full exchange into s-DDM^29^. CorA in sND was run on a Superose6 Increase 10/300 GL column (GE Healthcare) in 20 mM TrisDCl pH 7.5, 150 mM NaCl, 1 mM DTT in D_2_O with an initial flow rate of 0.5 ml/min. For all samples, the flow was reduced to 0.05 ml/min during elution of the main peak. Measurements were run twice to obtain data from two sample-to-detector distances, 2 m and 11 m, yielding data in a *q*-range of 0.0044 Å^-1^ to 0.42 Å^-1^ with the neutron wavelength of 6 Å. The scattering intensity, *I(q)*, was brought to absolute scale in units of cm^-1^ by normalizing to the direct beam intensity measured with an attenuator in place.

Pair distance, *p(r)*, distributions were calculated by Bayesian indirect Fourier transformation at the bayesapp server^32^ (available from the genapp server at https://somo.chem.utk.edu/bayesapp/). *p(r)*s and the corresponding *I(q)s* for PDB structures were calculated using CaPP (available at github.com/Niels-Bohr-Institute-XNS-StructBiophys/CaPP) with a water layer excess density of 6 % applied to the parts of the protein outside the membrane.

### Model building

For use in both SANS comparison, normal mode analysis, and simulations, we rebuilt the missing residues and sidechains of the closed PDB structure 4I0U. The 2-6 missing residues at each chain terminal was rebuilt using Modeller’s automodel functionality^49^. In the open cryo-EM structures, between 18-20 residues were missing in the N-terminal. These were rebuilt from the closed X-ray structure (4I0U) to allow for direct comparison to SANS measurements.

### Normal mode analysis

Non-linear normal mode analysis was performed using the NOn-Linear rigid Block NMA^50^ (NOLB) routine (available from https://hal.inria.fr/hal-01505843v2). The NOLB routine was integrated with PEPSI-SANS 2.2 (https://team.inria.fr/nano-d/software/pepsi-sans/) and a regularization algorithm to avoid unphysical structures. In the latter, *T = χ^2^ + αS* is minimized, where *S* constraints the secondary structure deviation. Scanning over different values of *α*, a plot of *χ^2^* vs. *S* is obtained. The best compromise between conservation of structure and the best fit to the data is chosen from the “elbow”-region (Figure 4-figure supplement 2).

### Metadynamics and reweighting

The simulation system was set up using the MARTINI3.0b coarse grained force field^51^ with elastic networks terms applied to the individual chains only with a 0.9 nm cut-off and a force constant of 500 kJ/(mol nm^2^). As MARTINI does not contain parameters for protein bound Mg^2+^ but rather models it as a free ligand with four waters bound, the Mg^2+^ ions bound in the structure were deleted. To avoid overly electrostatic repulsions from the remaining Mg^2+^ coordinating amino acids (Asp89, Asp179, Asp253 and Glu88), they were transformed into their protonated states. The POPC membrane was obtained from CHARMM-GUI^52,53^ using 450 POPCs in each bilayer, ensuring and the entire system was big enough for larger conformational changes. The system was solvated with the MARTINI non-polarizable water and neutralized with 300 mM NaCl comparable to the SANS experimental set-up. The equilibration was performed according to the CHARMM-GUI equilibration protocol using a minimization step followed by 6 equilibration steps with slow decrease in the positional restraint forces on both lipids and protein in each step^53^.

The GROMACS-5.1.4 software package^54^ was used to simulate with a 20 fs time step. Temperature and semi-isotropic pressure were controlled at 303.15 K and 1 bar using the stochastic velocity rescaling thermostat^55^ and Parrinello-Rahman barostat^56^. Electrostatic interactions were modulated using the reaction field approach. The cutoff of short-range distance for the electrostatic interactions was 1.1 nm. The potential shift Verlet scheme was used to cut off the Lennard-Jones potential at long ranges.

Well-Tempered Metadynamics^57^ simulations were performed with the PLUMED2.3.0 software^58^. A radius of gyration collective variable was applied on all back-bone beads for the intracellular residues 170 190 and 220 250 of all 5 chains. The metadynamics parameters were set as follows: Gaussian width = 0.05, Gaussian height = 2.1, Gaussian de-position stride = 100, biasfactor = 15 and an upper wall defined at a CV radius of gyration of 4.0 nm. The wall was defined as a harmonic restraint with a force constant = 10000, harmonic exponential power = 4, off-set = 0, and a rescaling factor of 1. Multiple wall types and sizes was attempted with lower walls causing too little dynamics for fitting with the experimental SANS data and higher walls causing individual monomers to bend unphysically and giving unfeasible large sample space. Clustering of the simulation trajectory was performed using the KMeans clustering method^59^.

The software program BME was used reweight the MetaD trajectory to fit the experimental SANS data^60^. The hyperparameter θ was determined based on a L-curve analysis (analogous to the procedure in normal mode analysis, see Figure 4-figure supplement 2) of S_rel_ vs χ^2^, where θ is chosen where a natural kink is observed and any further decrease in χ^2^ gives rise to an increasing larger penalty in S_rel_. As the simulations are not fully converged and the chosen force field is coarse grained, we set the trust in the force field lower and chose a slightly lower θ -value than the kink observed.

To account for a fixed fraction of symmetric pentamers, a differential intensity was derived by I_diff_ = I_exp_ – f · I_calc, 5sym_, where f is the fraction of symmetric pentamer, I_exp_ is the experimental SANS data, and I_calc,5sym_ is the SANS signal of PDB 4I0U calculated by PEPSI-SANS. The forward scattering of the calculated SANS signal, I_calc,5sym_(0), was scaled to match the forward scattering of the experimental SANS data. Reweighting was done against the differential SANS signal.

### Solid-state NMR Spectroscopy

Spectra of uniformly labelled samples were measured at a magnetic field of either 19.5 T or 23.4 T corresponding to a ^1^H Larmor frequency of 800 MHz and 1000 MHz, respectively. The spectrometers were equipped with a Bruker 0.7 mm MAS probe, spinning at 107 kHz, at a constant temperature of 300 K. Spectra of ^15^N Ala labelled samples were acquired at 23.4 T on a Bruker 1.3 mm MAS probe, spinning at 60 kHz, at a constant temperature of 300 K. The assignment of backbone resonances was preliminary obtained by acquiring a set of ^1^H-detected 3D experiments as described in ^61,62^. Adamantane was used as the external reference. Spectral analysis and assignment were accomplished with CcpNmr Analysis ^63^ and FLYA ^64^.

Relaxation experiments were based on a ^1^H,^15^N ^1^H-detected CP-HSQC experiment incorporating an appropriate relaxation delay^65^. 38 and 33 residues spanning different regions of the proteins were used for the uniformly labelled sample in the presence and in the absence of Mg^2+^, respectively. The measurements of site-specific ^15^N R_1ρ_ rates were performed at 107 kHz MAS, 19.5 T and 300 K and 280 K, using relaxation delays of 0.05, 1, 5, 15, 50, 100, 200 ms under a spin-lock field of 15 kHz. Measurements of bulk ^15^N R_1_ were performed at 107 kHz MAS, 19.5 T and 300 K, using relaxation delays of 0.5, 1, 2.8, 6.8, 15.8, 23.8, 53.5, 80 s. Measurements of ^15^N R_1ρ_ rates were additionally performed on the ^15^N Ala labelled samples in the presence and in the absence of Mg^2+^ at 60 kHz MAS, 23.5 T and 300 K using relaxation delays of 0.1, 1, 5, 10, 25, 50, 100 ms under a spin-lock field of 15 kHz. The relaxation rates were obtained by fitting the experimental decay curves with a mono-exponential function. The error was estimated from the experimental noise by use of a Monte-Carlo evaluation.

### Activity assay

Large unilamellar vesicles of POPC were prepared by dissolving a lipid film in 10 mM MOPS-KOH pH 7.2, 150 mM KCl, 100 μM EGTA including 10 µM Mag-Fluo-4 (Thermo) to a POPC concentration of 15 mg/ml, which was extruded through 0.2 µm membrane filters for 35 times using a mini-extruder (Avanti Polar Lipids). CorA was inserted by mixing a sample of 10.5 mg/ml LUVs, 2 µM CorA, 10 µM Mag-Fluo-4 and 50 mM octyl glucoside. Biobeads SM-2 were added to 45 % w/v and incubated at RT for 30 min, before purifying and at the same time exchanging the extravesicular buffer to measurement buffer (10 mM MOPS-KOH pH 7.2, 150 N-methyl-D-glucamine-HCl, 100 μM EGTA) on Sephadex G50 resin. 20 µl of CorA-LUVs (or plain LUVs) were diluted to a total of 1 ml in measurement buffer (prepared in H_2_O or D_2_O, respectively) containing 10 µM valinomycin with or without 1 mM Co[NH_3_]^3+^ present. CorA activity was monitored by Mag-Fluo-4 fluorescence at 515 nm (excitation at 488 nm) on a FluoroMax fluorometer (Horiba) upon addition of 10 mM MgCl_2_ (from a 1 M stock prepared in H_2_O or D_2_O, respectively). The signal was normalized to the flat signal recorded before addition of MgCl_2_.

### Negative stain EM

CorA was purified by SEC and diluted to 0.1 μM in appropriate buffers containing 1 mM EDTA or 40 mM Mg^2+^. Copper grids were neutralized with an Easiglow glow discharger (Agar Scientific). 3 μl of sample was applied to the grid and incubated for 30 s. The grid was blotted onto a filter paper from the edge, and 3 μL of 2% uranyal formate was added immediately and incubated for 30 s. The staining procedure was repeated two more times. After the final staining, the grid was left to dry for ten minutes. EM data were acquired on a Tecnai TEM (FEI, Thermo Fischer scientific) at Aarhus University, Denmark. The micrographs were processed by XMIPP to *.mcp files, and particle picking, 2D class averages and 3D model refinement was done in Relion 3.0. Statistics for the 3D refinement are given in Table 1.

**Table 1.**
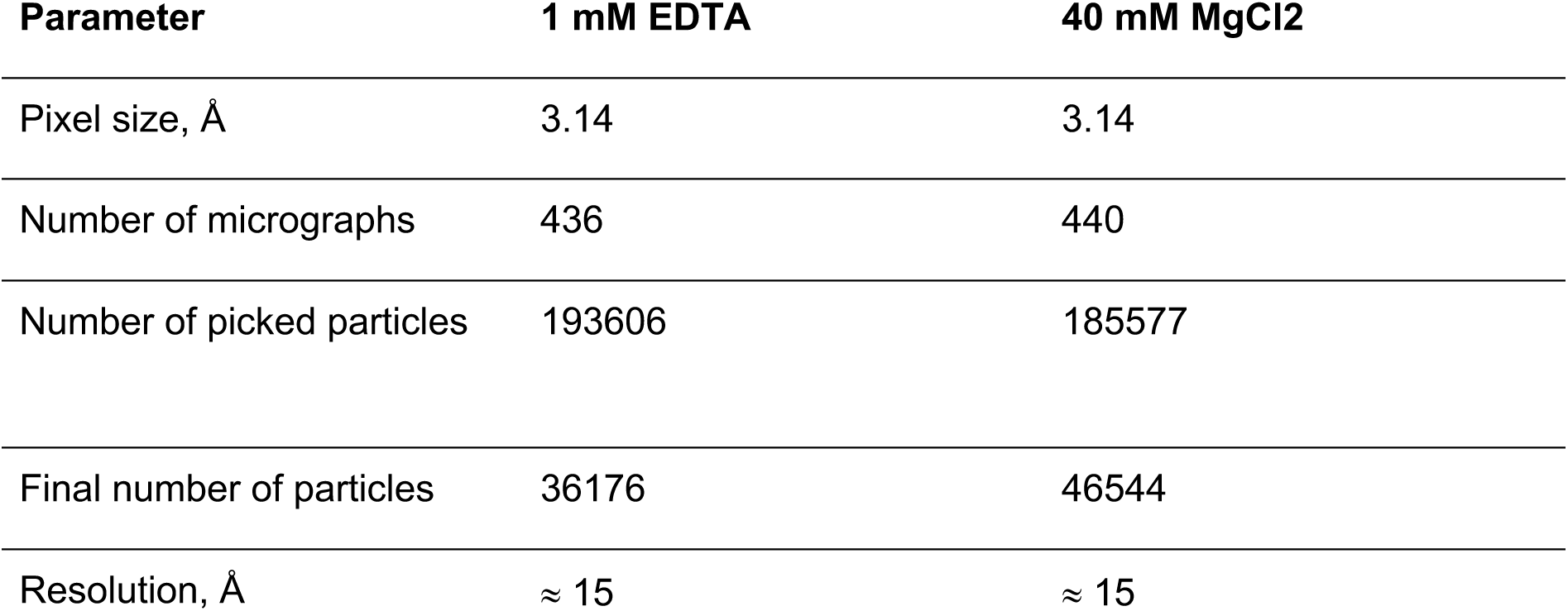
Negative stain EM statistics for 3D model refinement.

| Parameter | 1 mM EDTA | 40 mM $\text{MgCl}_2$ |
| --- | --- | --- |
| Pixel size, $\text{\AA}$ | 3.14 | 3.14 |
| Number of micrographs | 436 | 440 |
| Number of picked particles | 193606 | 185577 |
| Final number of particles | 36176 | 46544 |
| Resolution, $\text{\AA}$ | $\approx 15$ | $\approx 15$ |

## Acknowledgements

We thank Elliot Gilbert for his assistance with SANS experiments at QUOKKA at ANSTO and Marta Brennich for her assistance with SAXS experiments at BM29 at the ESRF. Thomas Boesen is acknowledged for his help with EM experiments conducted at Aarhus University and Michael Gajhede for his assistance in EM data processing. We thank Ida Louise Jørgensen for helping with functional reconstitution of CorA in large unilamellar vesicles and Michael Maguire for providing a plasmid encoding CorA.

## Additional information

### Funding

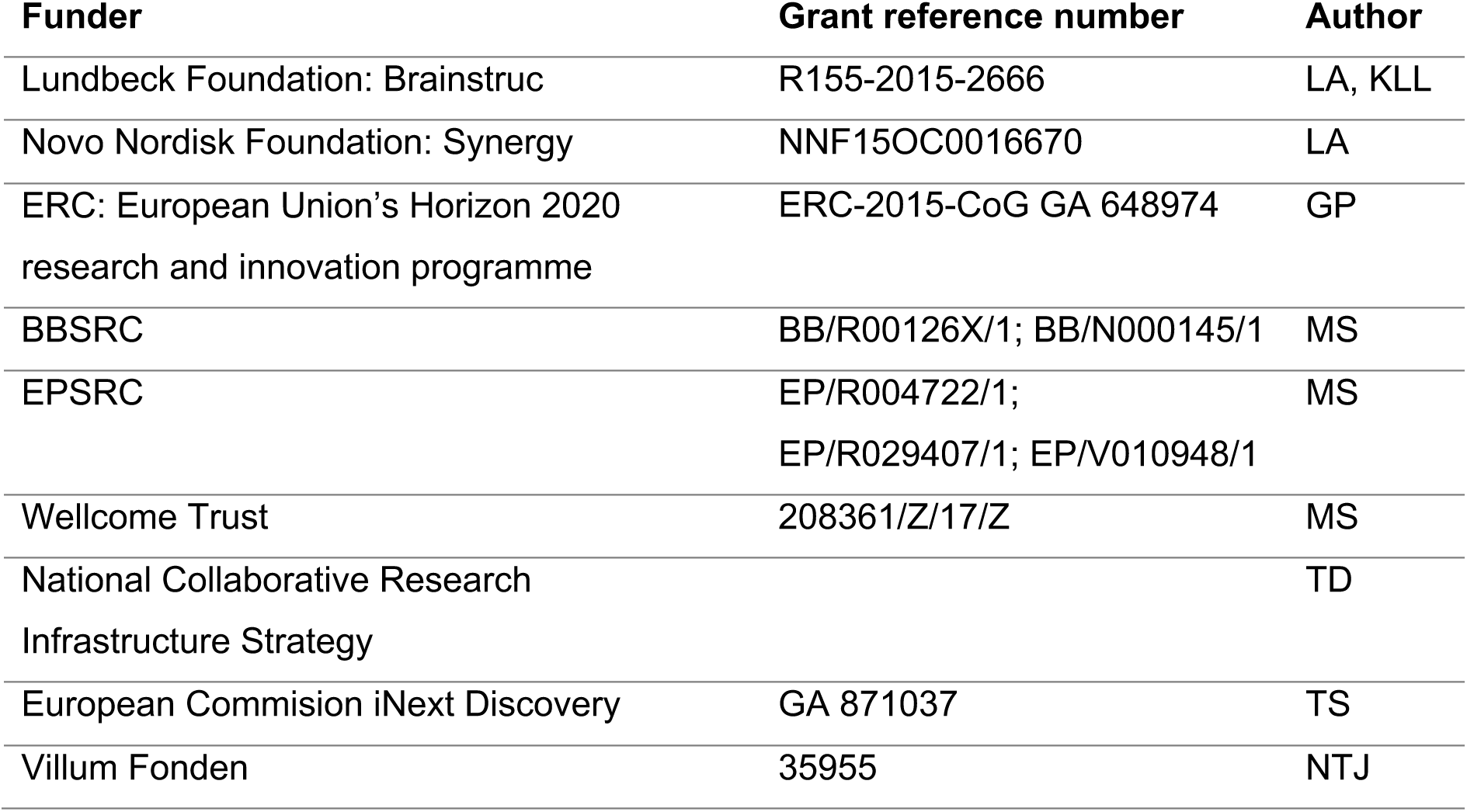

### Author Contributions

LA, GP and KLL conceived the project. NTJ, JB, NY, and TS prepared samples. NTJ, AHL, FGT, PH, AM, and MCP carried out SANS experiments and modeling. NTJ and TGP carried out activity measurements. NTJ and AHL did EM experiments. TB, AHL, and RC carried out simulations. MB, AB, and TS carried out NMR experiments and modeling. LA, GP, KLL, MR, TGP, TD, and MS supervised the research. NTJ, MB, TS, LA, GP and KLL wrote the manuscript with input from all authors.

### Data availability

The SANS data have been deposited to the Small Angle Scattering Biological Data Bank (https://www.sasbdb.org/, access codes SASDM42, SASDM52, SASDM62, SASDM72). The NMR resonance assignments have been deposited in the Biological Magnetic Resonance Data Bank (https://bmrb.io, access code 50959). The EM maps have been deposited to the Electron Microscopy Data Bank (https://www.ebi.ac.uk/emdb, access codes EMD-13326 and EMD-13327). Fluorescence data on CorA activity are available at GitHub https://github.com/Niels-Bohr-Institute-XNS-StructBiophys/CorAData/. The metadynamics simulations are available at GitHub https://github.com/KULL-Centre/papers/tree/main/2021/CorA-Johansen-et-al.

---

**Figure 2-figure supplement 1.**
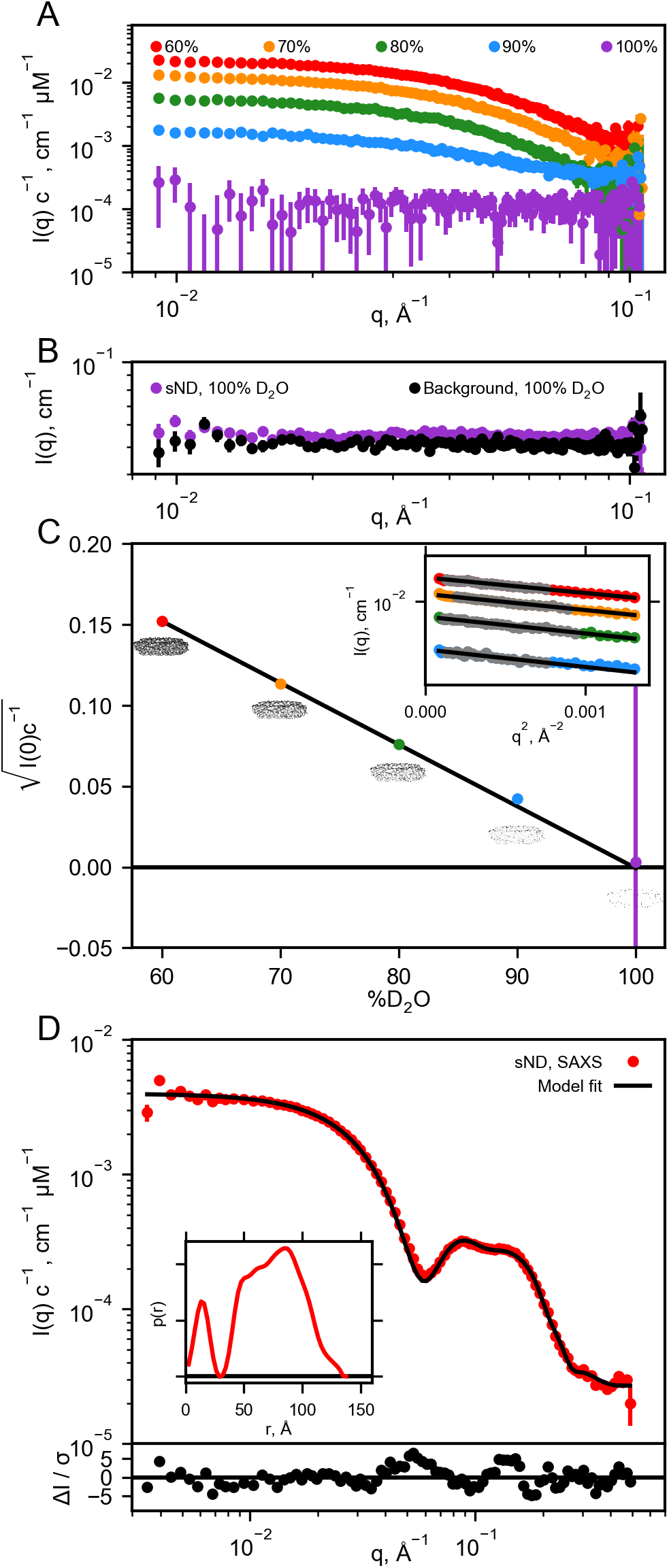
Validation of nanodisc match-out deuteration. A: Background subtracted SANS data measured on sNDs in different percentage of D_2_O. B: Comparison of unsubtracted SANS data and corresponding background measurement in 100 % D_2_O. C: Plot of contrast vs D_2_O to verify match-point of sample (fitted to 99.9 % D_2_O by linear regression). I(0) values were determined using Guinier fits to the low-q data (grey points) as highlighted in the insert. The Guinier fit to the SANS data at 100 % D_2_O has not been shown due to the large errorbars, and the fit was also not included for fitting the contrast vs D_2_O. D: SAXS data on sND collected in H_2_O buffer together with p(r) distribution (insert). A nanodisc model^33^ was fitted to the data using WillItFit^34^, producing a great fit, as evaluated by the residual plot on the bottom.

**Figure 2-figure supplement 2.**
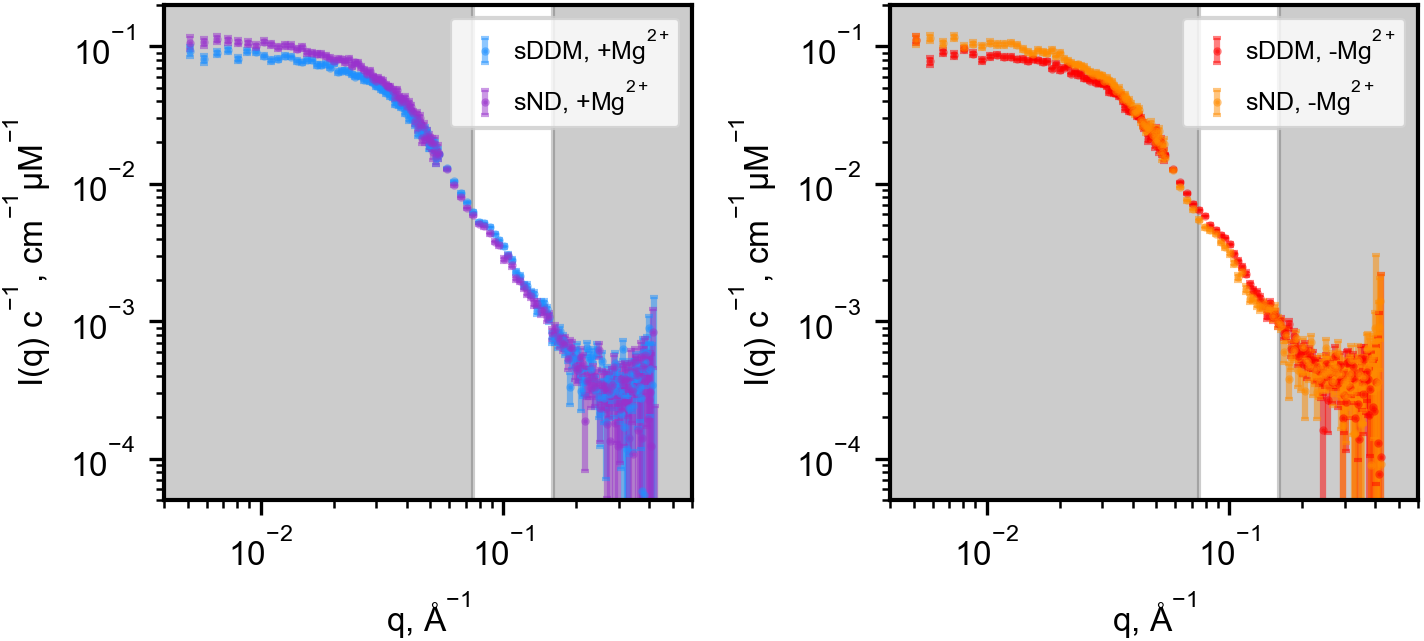
Comparison of SANS data in sDDM and sND. Same data as shown in Figure 1D-E, but presented here to compare data collected in sDDM vs sND. Despite the slightly higher forward scattering in the sND data, the SANS data from the two different systems overlap in the region of interest, as highlighted. Given that CorA had spent less time and steps in presence of DDM prior to reconstitution in sNDs, the excess signal likely stems from a small number of tightly bound E. coli lipids that do not match out in D_2_O. By comparing the extrapolated values of I(0) between the sDDM and sND samples, we estimated the number to be on the order of 10 lipids, i.e. two lipids per protomer. Although the presence of visible lipids would complicate further detailed analysis, the striking identicality of the measured curves with and without Mg^2+^ in sNDs supports no large structural rearrangements.

**Figure 4-figure supplement 1.**
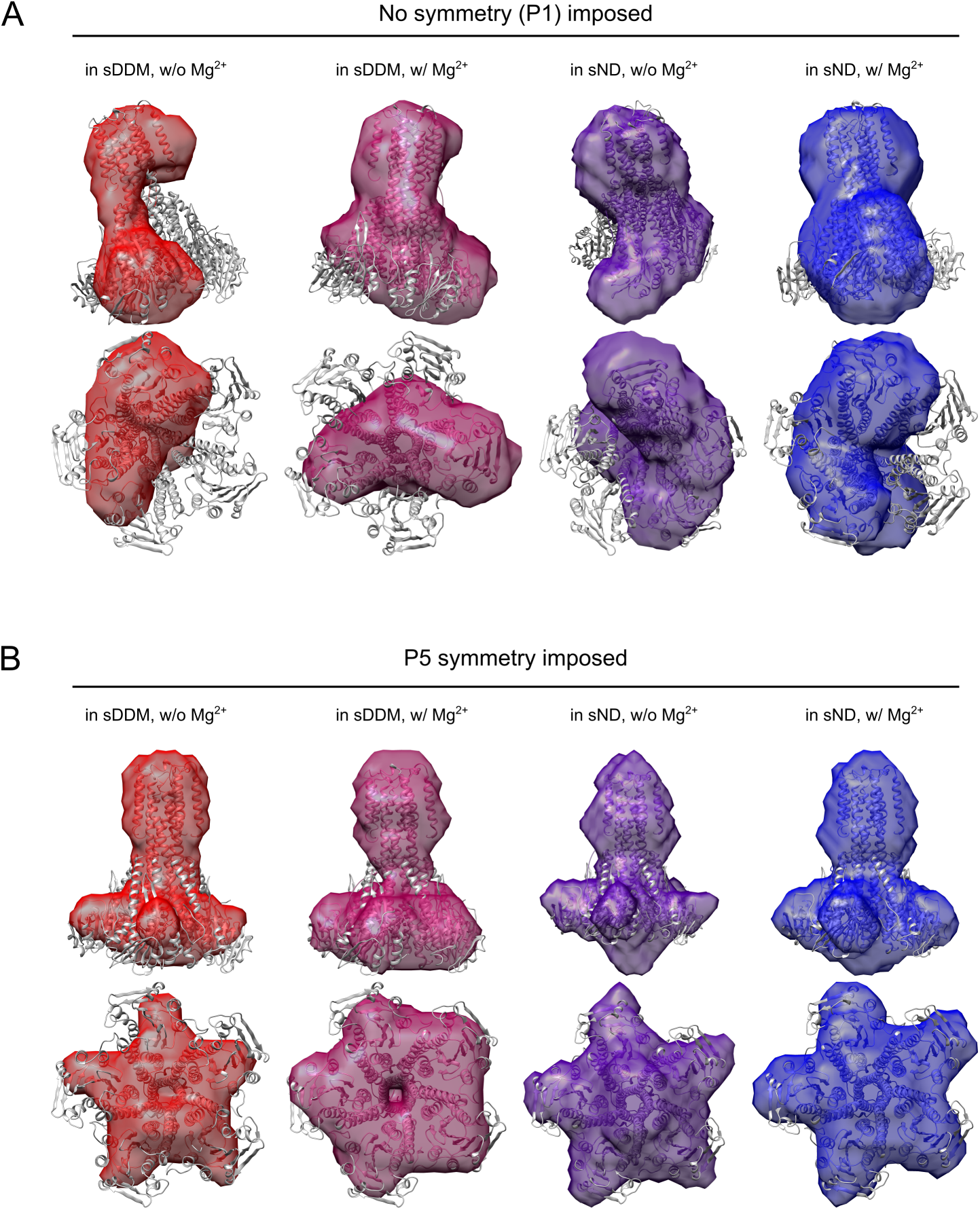
Bead modeling of CorA. All four SANS data sets were analysed by the DAMMIF pipeline^38^ available from the ATSAS package^39^. In total, DAMMIF was run 100 times in slow mode with either P1 (A) or P5 (B) symmetry selected. For P1 symmetry, the normalized spatial discrepancy (NSD) values varied from 1.2 to 1.5, whereas for P5 symmetry, NSD values varied from 1.4 to 1.7. NSD values above 1 indicate that the calculated models differ systematically from one another.

**Figure 4-figure supplement 2.**
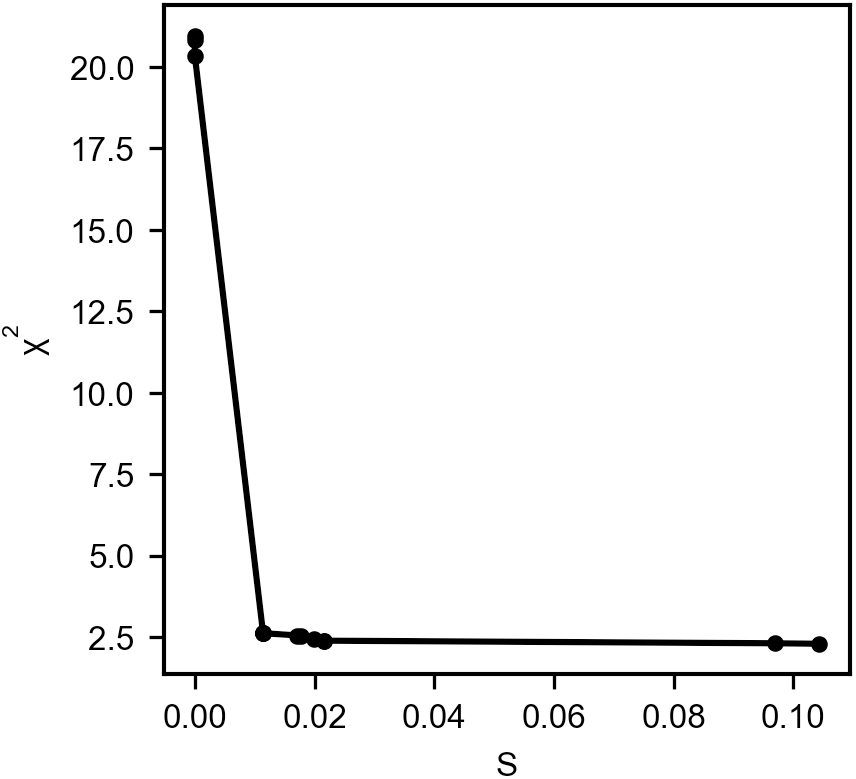
Goodness of fit from NMA generated structures to SANS. The best compromise between conservation of structure and the best fit to the data is in the “elbow”-region, here chosen at S = 0.01 (with α = 600).

**Figure 4-figure supplement 3.**
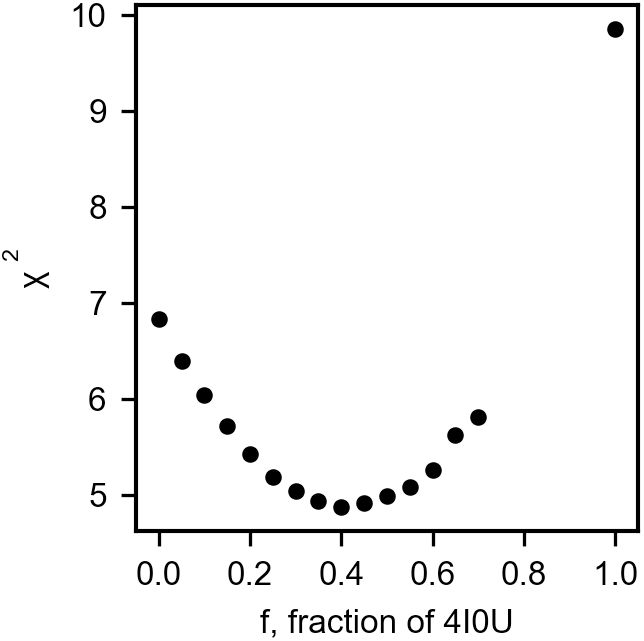
Ensemble fitting to SANS data with symmetric CorA included. The goodness of fit (χ^2^) plotted as a function f, the fraction of 4I0U, fixed before BME reweighting of the MetaD generated ensemble to fit the SANS data. The data on the graph were fitted by a quadratic function to yield the minimum with standard deviation at 40 ± 28 % 4I0U, where the standard deviation was calculated by an increment in χ^2^ of 1.

**Figure 5-figure supplement 1.**
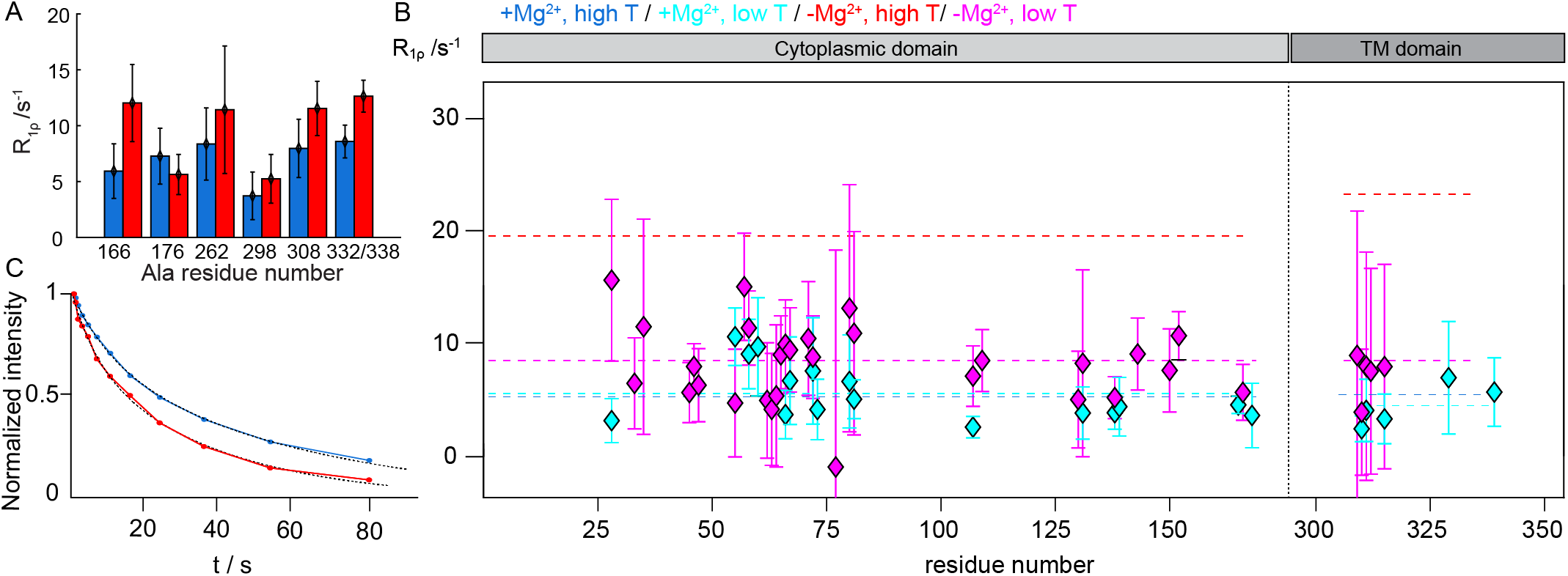
CorA backbone dynamics by MAS NMR: Site-specificity and temperature dependence. A: Site-specific ^15^N R_1ρ_ rates measured with a 15 kHz spin-lock field in the ^15^N-Ala sample of CorA. B: Comparison of site-specific ^15^N R_1ρ_ rates measured with a 15 kHz spin-lock field at ∼280 K in the presence (cyan) and in the absence (magenta) of Mg^2+^. Dashed lines indicate the average relaxation rate values in the ICD and TMD distinctly, at higher (blue and red ) and lower (cyan and magenta) temperature. C: Bulk ^15^N R_1_ decay curves measured at 300 K. Experimental curves were fitted with a biexponential decay.

**Figure supplement 1.**
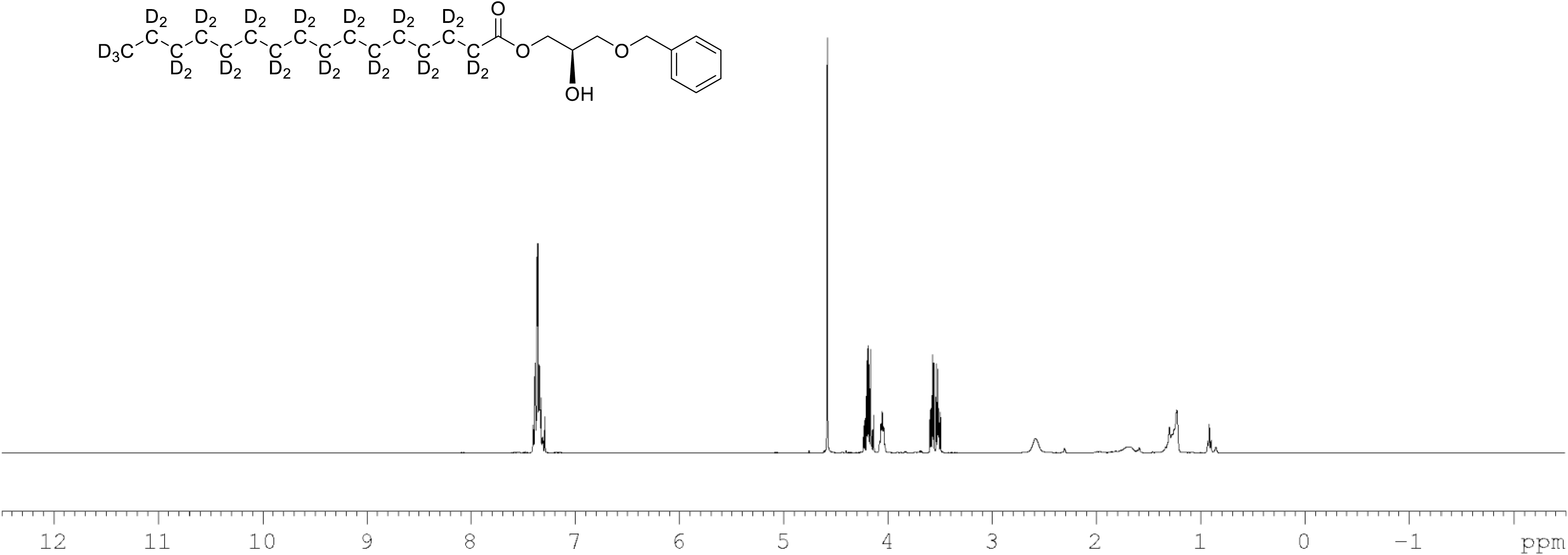
^1^H NMR of 1-palmitoyl-d_31_-sn-3-benzyloxy-glycerol (Figure 7, molecule **2**) in CDCl_3_. (400 MHz, CDCl_3_), δ residual protons 0.88 (m, 0.22H), 1.10-17 (m, 1.96H), 1.57.1.86 (m, 1.55H), 1.96 (m, 0.54H), 3.69 (m, 2H), 4.09 (m, 1H), 4.22 (m, 2H) 4.60 (s, 2H), 7.36 (m, 5H).

**Figure supplement 2.**
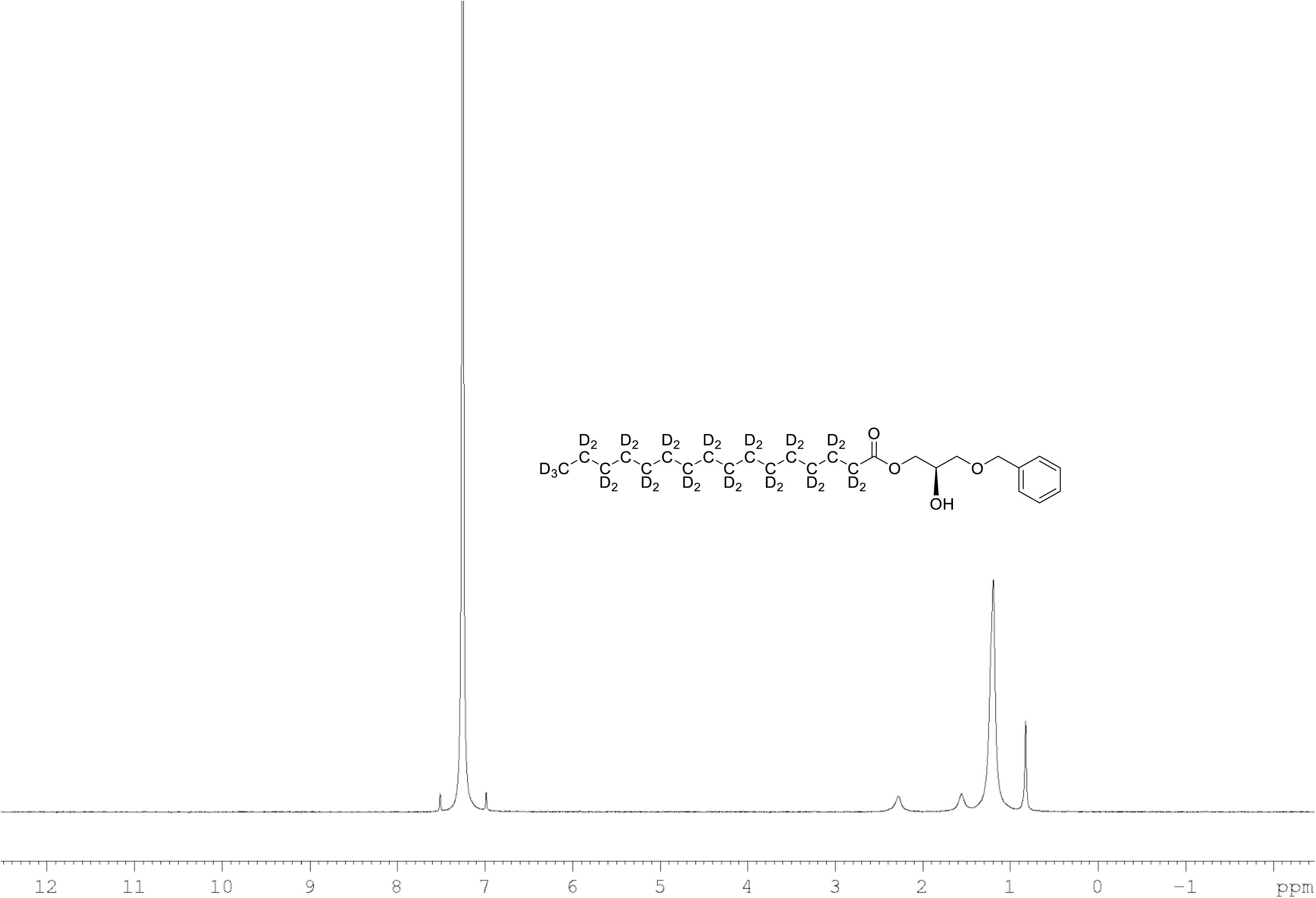
^2^H NMR of 1-palmitoyl-d_31_-sn-3-benzyloxy-glycerol (Figure 7, molecule **2**) in CDCl_3_. (400 MHz, CDCl_3_), δ 0.82 (m), 1.18 (m), 1.54 (m), 2.27 (m).

**Figure supplement 3.**
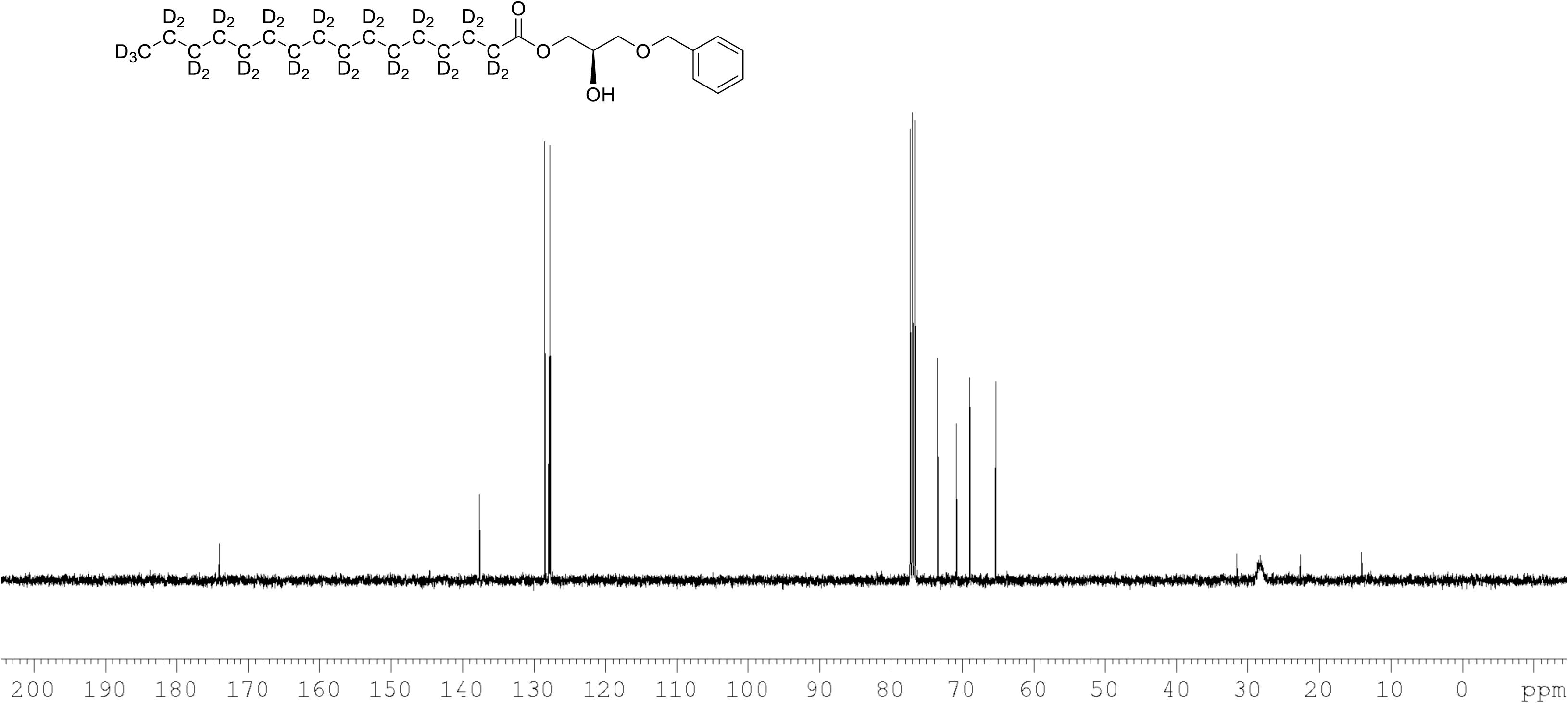
^13^C NMR of 1-palmitoyl-d_31_-sn-3-benzyloxy-glycerol (Figure 7, molecule **2**) in CDCl_3_. (400 MHz, CDCl_3_), 10.9 (m), 22.09 (m), 28.33 (m), 33.05 (m), 65.33, 68.9, 70.80, 73.5, 127.7, 127.9, 128.5, 137.6, 174.0.

**Figure supplement 4.**
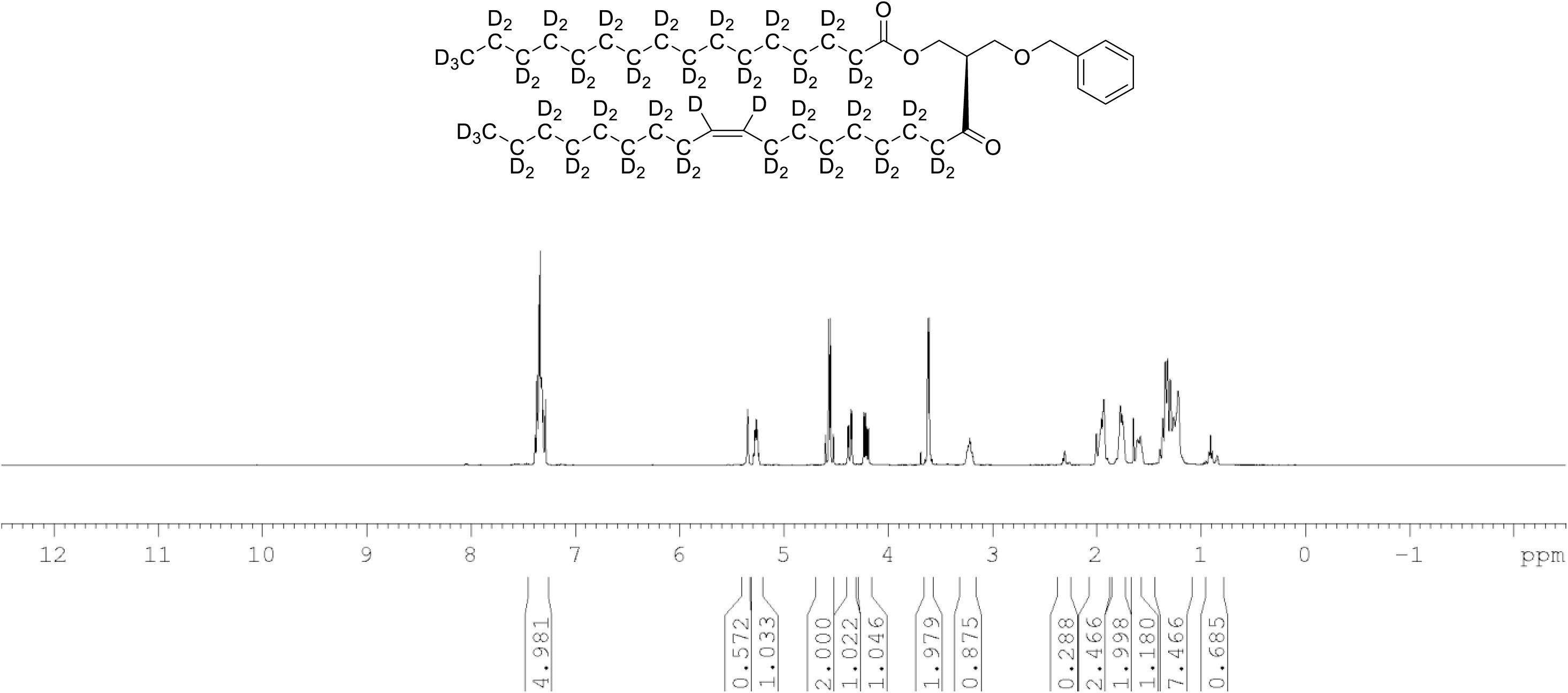
^1^H NMR of 1-palmitoyl-d_31_-2-oleoyl-d_33_-sn-3-benzyloxy-glycerol (Figure 7, molecule **3**) in CDCl_3_. (400 MHz, CDCl_3_), δ residual protons 0.90 (m, 3.56H), 1.29 (m, 5.15H), 1.98 (m, 0.68H), 2.29 (m, 1.25H), 3.61 (d, J = 5.0Hz, 2H), 4.21 (m, 1H), 4.36 (m, 1H), protonated benzyl protons 4.56 (AB q, J = 12Hz, 2H), 5.26 (m, 1H), 5.35 (s, 0.65H), 7.34 (m, 5H).

**Figure supplement 5.**
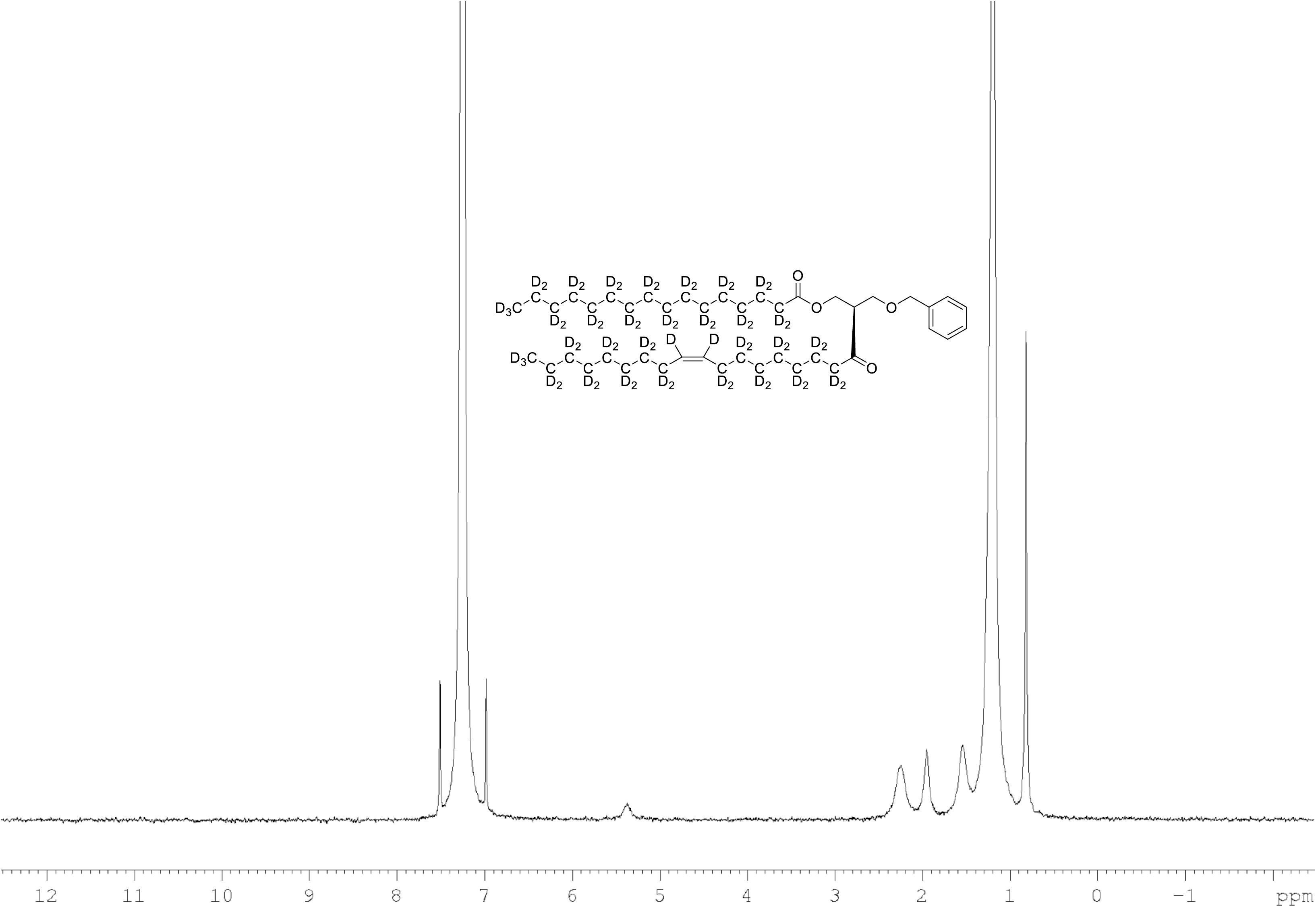
^2^H NMR of 1-palmitoyl-d_31_-2-oleoyl-d_33_-sn-3-benzyloxy-glycerol (Figure 7, molecule **3**) in CDCl_3_. (400 MHz, CDCl_3_), δ 0.82 (m, 6D), 1.19 (m, 35.35D), 1.53 (m, 3.74D), 1.94 (m, 3.34D), 2.25 (m, 2.63D), 3.56 (m, 1.71D), 4.27 (m, 1.04D), 5.35 (m, 1.92)

**Figure supplement 6.**
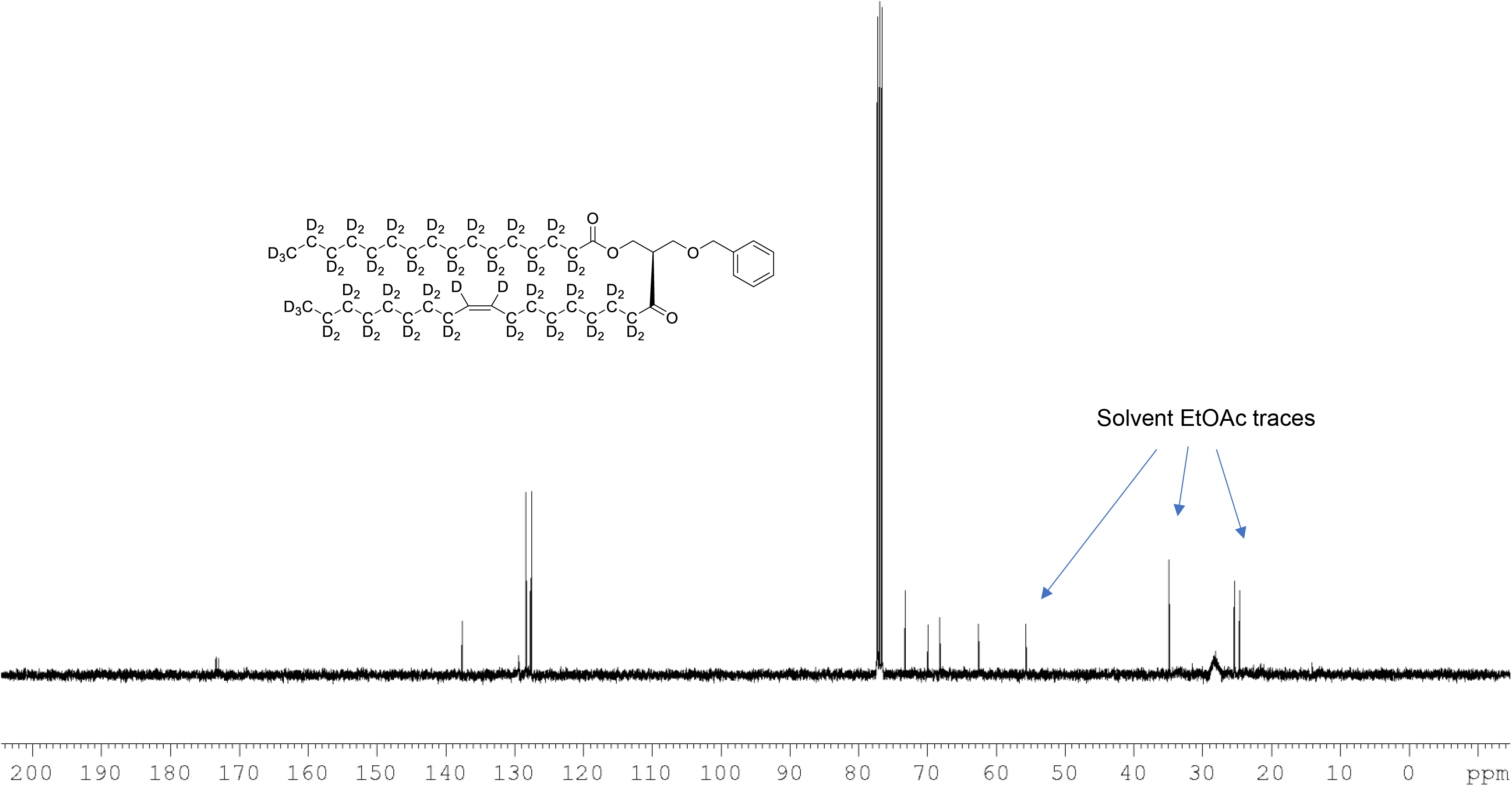
^13^C NMR of 1-palmitoyl-d_31_-2-oleoyl-d_33_-sn-3-benzyloxy-glycerol (Figure 7, molecule **3**) in CDCl_3_. (400 MHz, CDCl_3_), δ 13.08 (m), 21.60 (m), 23.96 (m), 26.50 (m), 28.2 (m), 30.60 (m), 33.80 (m), 62.60, 68.39 9, 70.10, 73.1, 127.7, 127.9, 128.5, 137.7, 173.1, 173.4.

**Figure supplement 7.**
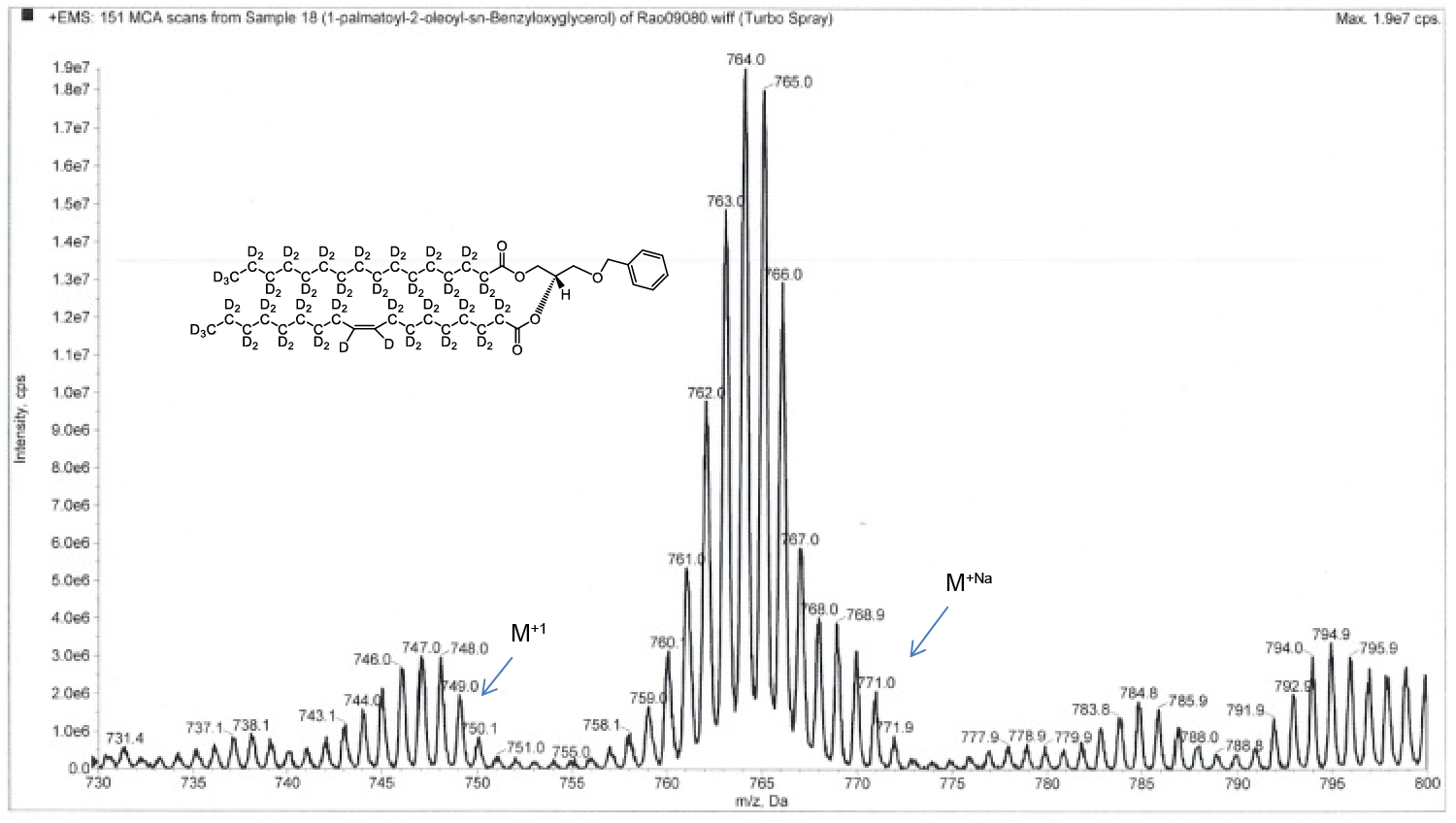
ESI-MS, m/z 749 [M^+1^]^+^ of POPC precursor 1-palmtoyl-2-oleoyl-sn-benzyloxyglycerol-d_64_ (Figure 7, molecule **3**). Overall 94%D, isotope distribution d_64,_ 7.8%, d_63,_ 15.1%, d_62,_ 19.5%, d_61,_ 18.0%, d_60,_ 13.1%, d_59,_ 10.2%, d_58,_ 6.5%, d_57,_ 3.9%, d_56,_ 2.7%.

**Figure supplement 8.**
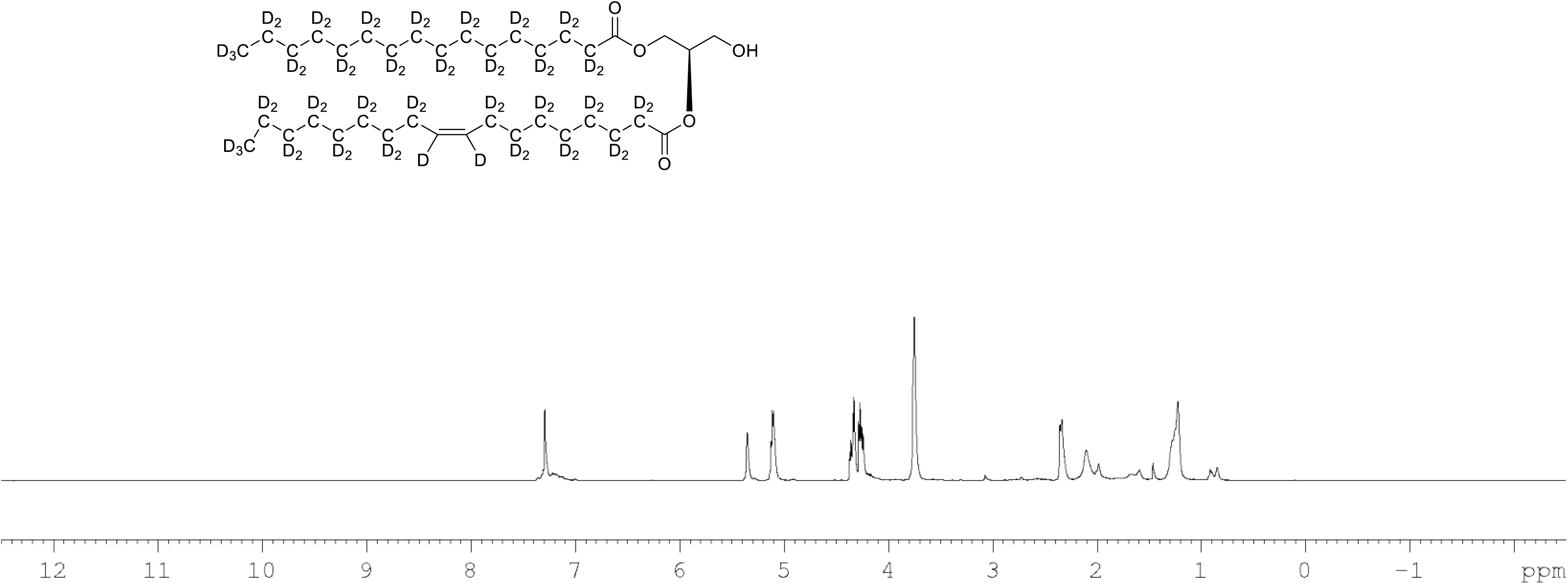
^1^H NMR of 1-palmitoyl-d_31_-2-oleoyl-d_33_-sn-3-glycerol (Figure 7, molecule **4**) in CDCl_3_. (400 MHz, CDCl_3_), δ residual protons 0.24 (m, 0.47H), 1.28 (m, 1.51H), 1.67 (m, 2.24H), 2.13 (m, 0.52H), 2.23 (m, 1.29H), 3.72 (m, 2H), 4.30 (m, 2H), 5.10 (m, 1H), 5.34 (s, 0.48H).

**Figure supplement 9.**
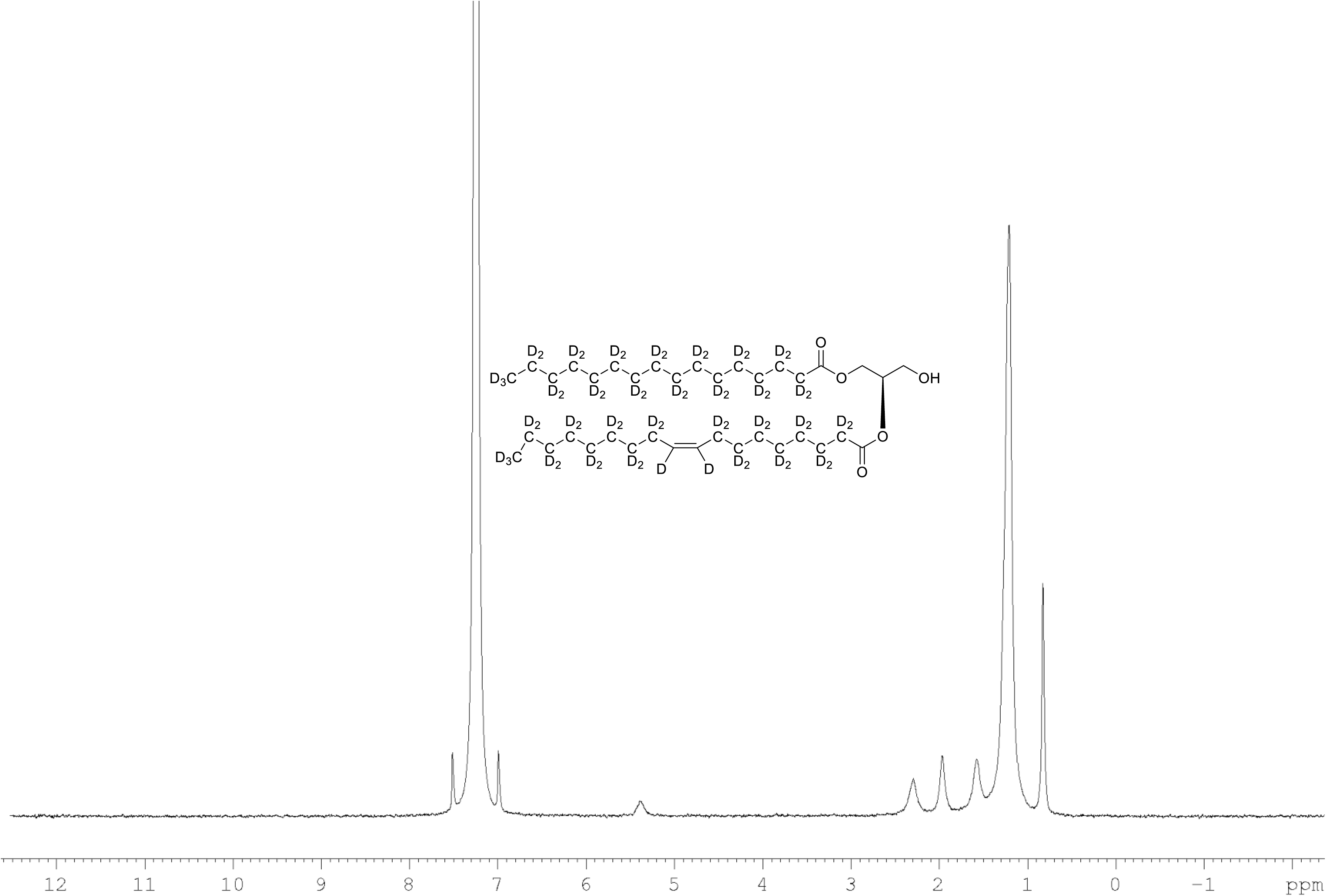
^2^H NMR of 1-palmitoyl-d_31_-2-oleoyl-d_33_-sn-3-glycerol (Figure 7, molecule **4**) in CDCl_3_. (400 MHz, CDCl_3_), δ 0.82 (m, 8.6D), 1.19 (m, 49.2D), 1.55 (m, 4.8D), 1.94 (m, 3.46D), 2.28 (m, 3.82D), 5.38 (m, 1.1D).

**Figure supplement 10.**
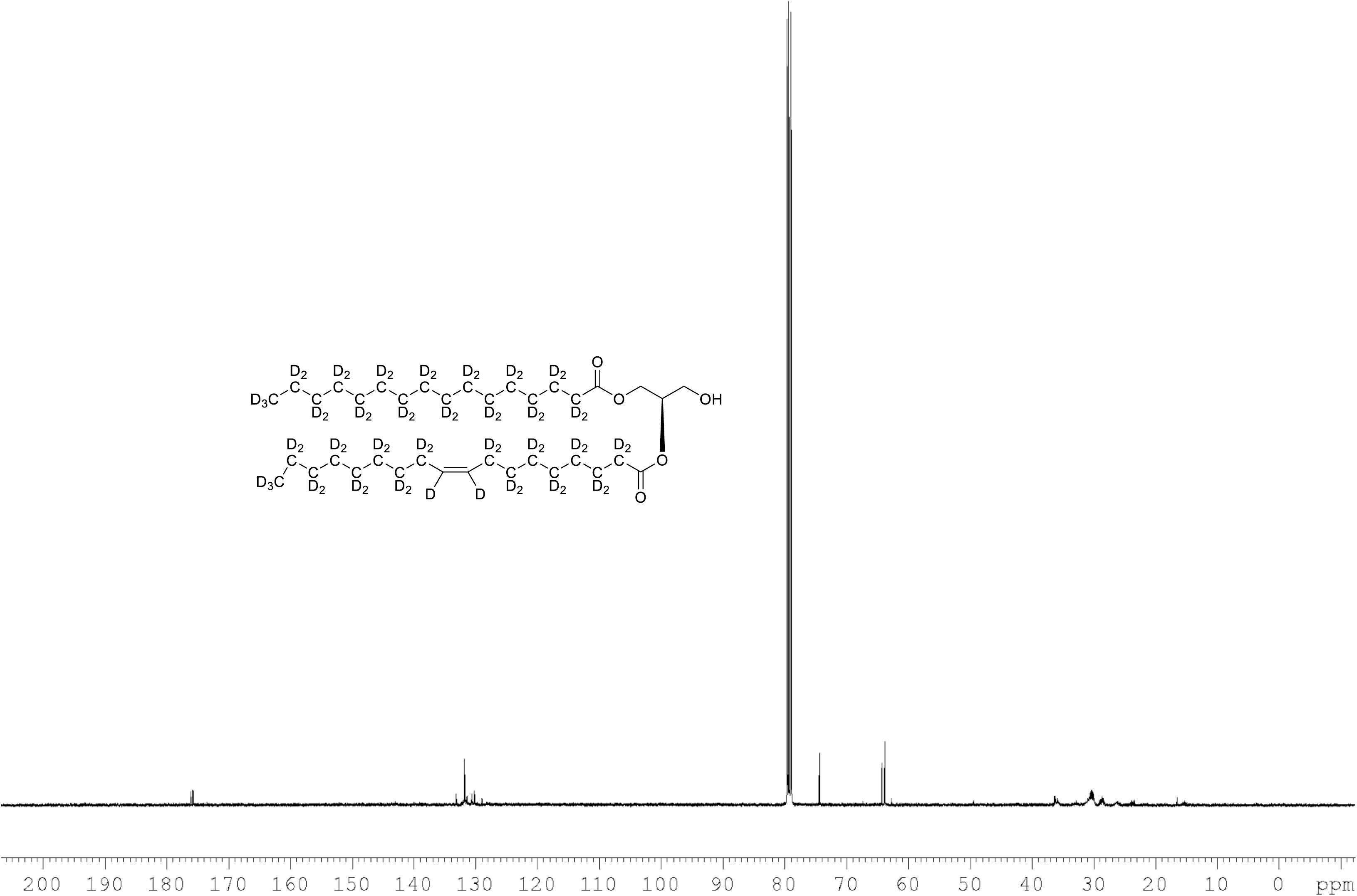
^13^C NMR of 1-palmitoyl-d_31_-2-oleoyl-d_33_-sn-3-glycerol (Figure 7, molecule **4**) in CDCl_3_. (400 MHz, CDCl_3_), δ 12.89 (m), 21.39 (m), 22.6 (m), 24.00 (m), 26.25 (m), 28.2 (m), 30.49 (m), 33.86 (m), 65.00, 60.76, 61.60, 71.64, 128.02, 129.14, 129.5, 173.53, 173.93.

**Figure supplement 11.**
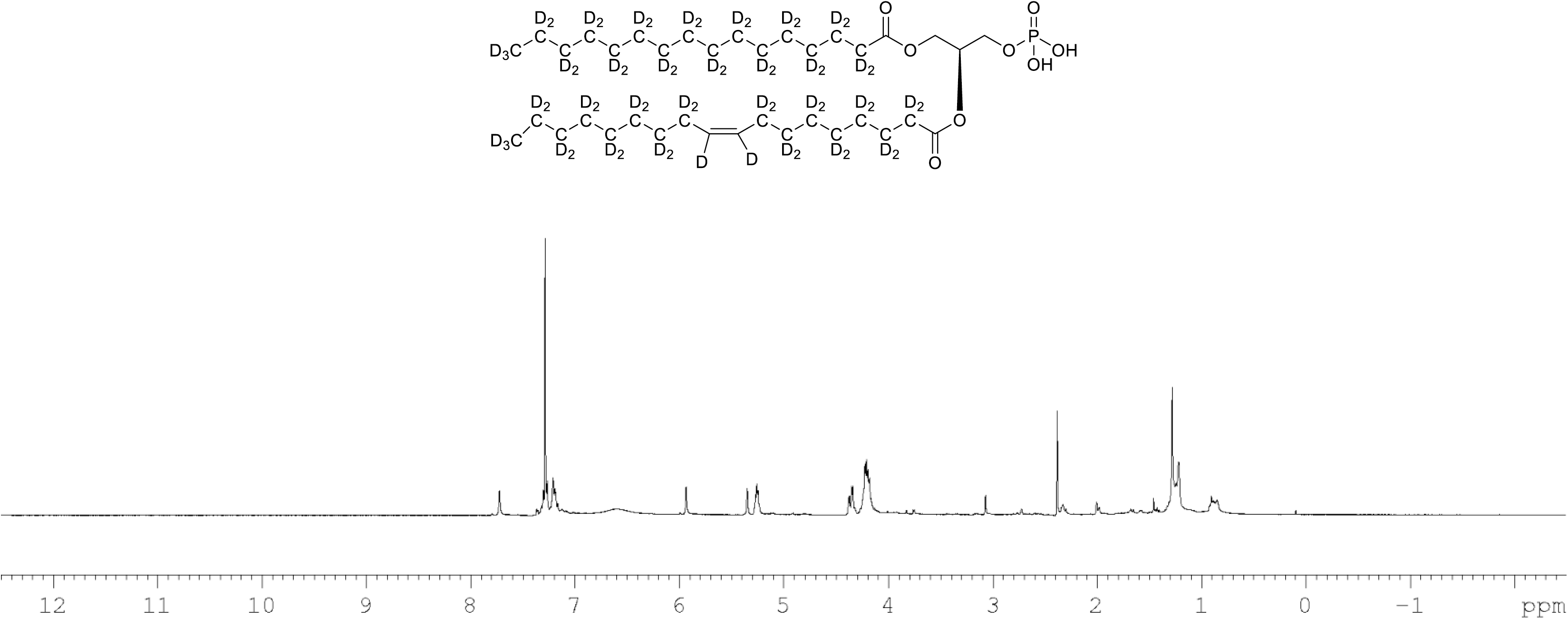
^1^H NMR of crude product 1-palmitoyl-d_31_-2-oleoyl-d_33_-sn-3-glycero-phosphatidic acid (Figure 7, molecule **5**) in CDCl_3_. This synthesised crude dried product was used in next step without further purification.

**Figure supplement 12.**
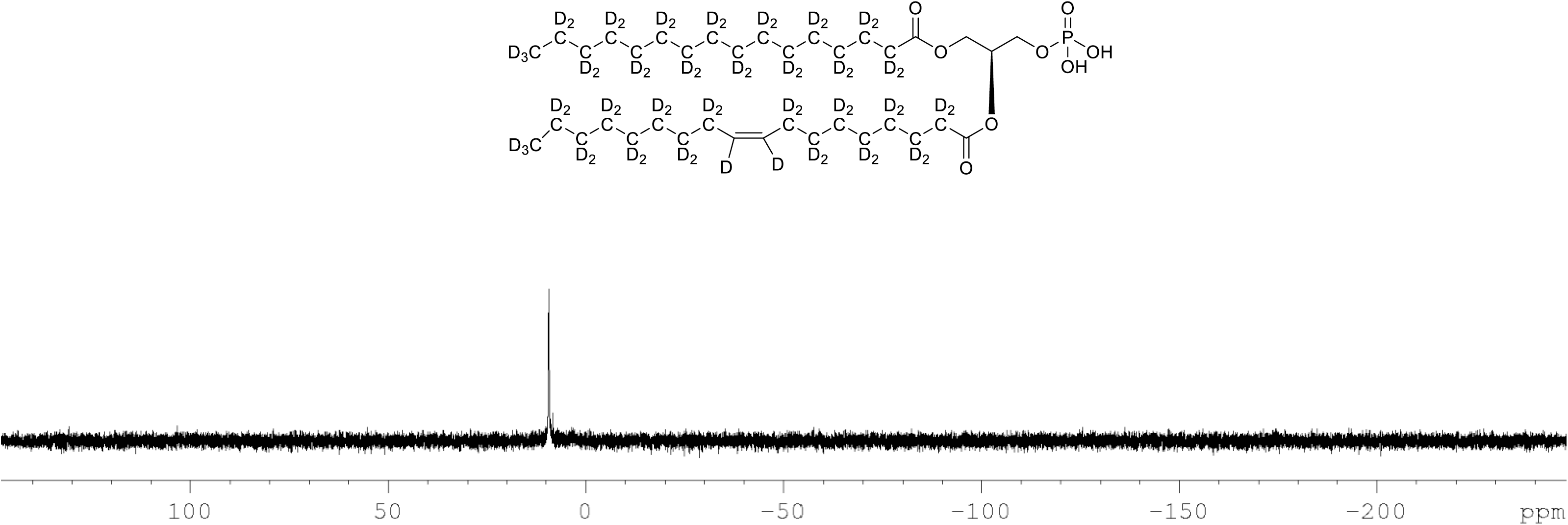
^31^P NMR of crude product 1-palmitoyl-d_31_-2-oleoyl-d_33_-sn-3-glycero-phosphatidic acid (Figure 7, molecule **5**) in CDCl_3_. This synthesised crude dried product was used in next step without further purification.

**Figure supplement 13.**
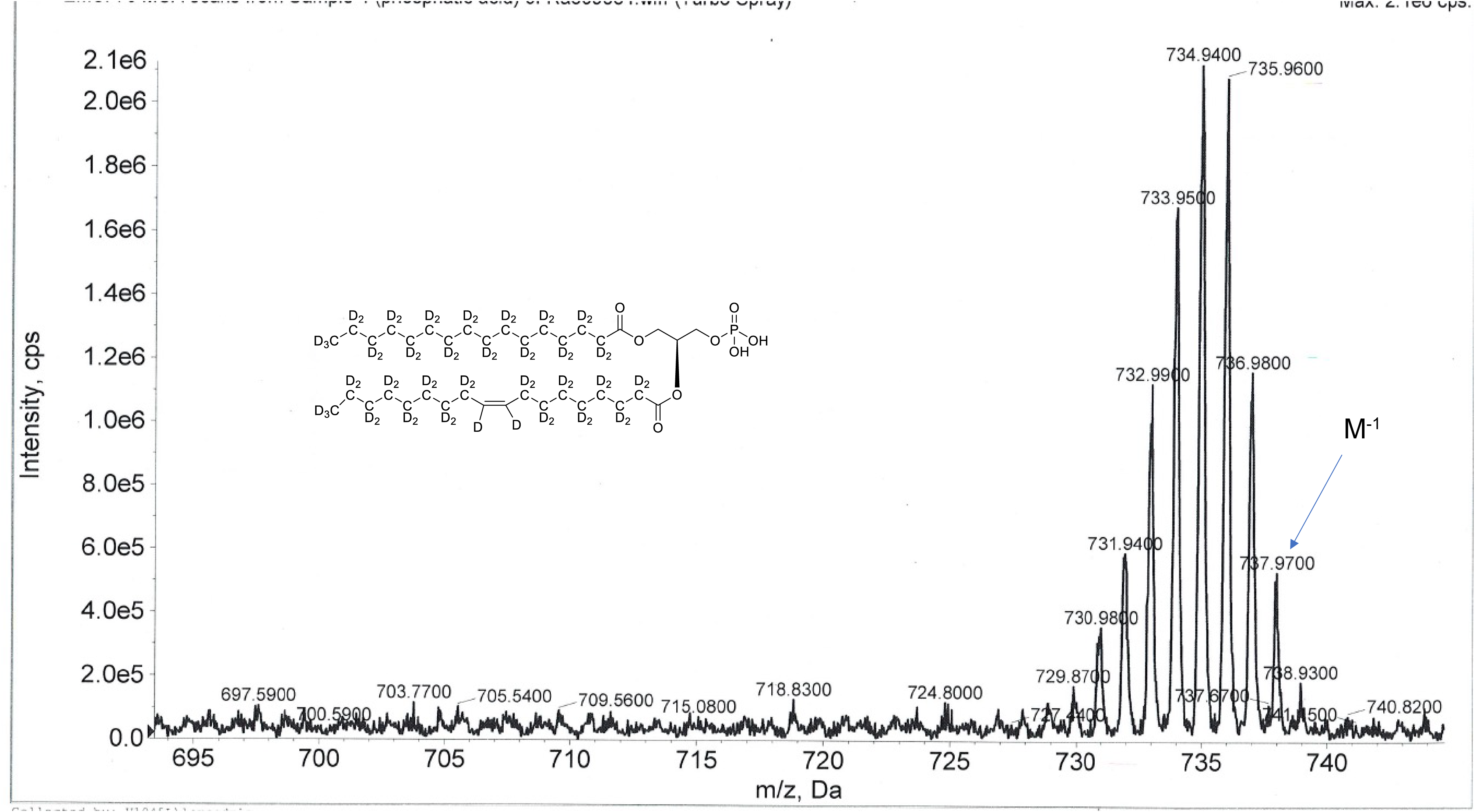
ESI-MS, m/z 737 [M^-1^]^-^ of crude product of 1-palmitoyl-d_31_-2-oleoyl-d_33_-sn-3-glycero-phosphatidic acid (Figure 7, molecule **5**). Overall 94%D, isotope distribution d_64,_ 0.6%, d_63,_ 8.9%, d_62,_ 18.8%, d_61,_ 25.1%, d_60,_ 20.6%, d_59,_ 13.0%, d_58,_ 7.5%, d_57,_ 5.2%, d_56,_ 0.2%, d_55,_ 0.1%.

**Figure supplement 14.**
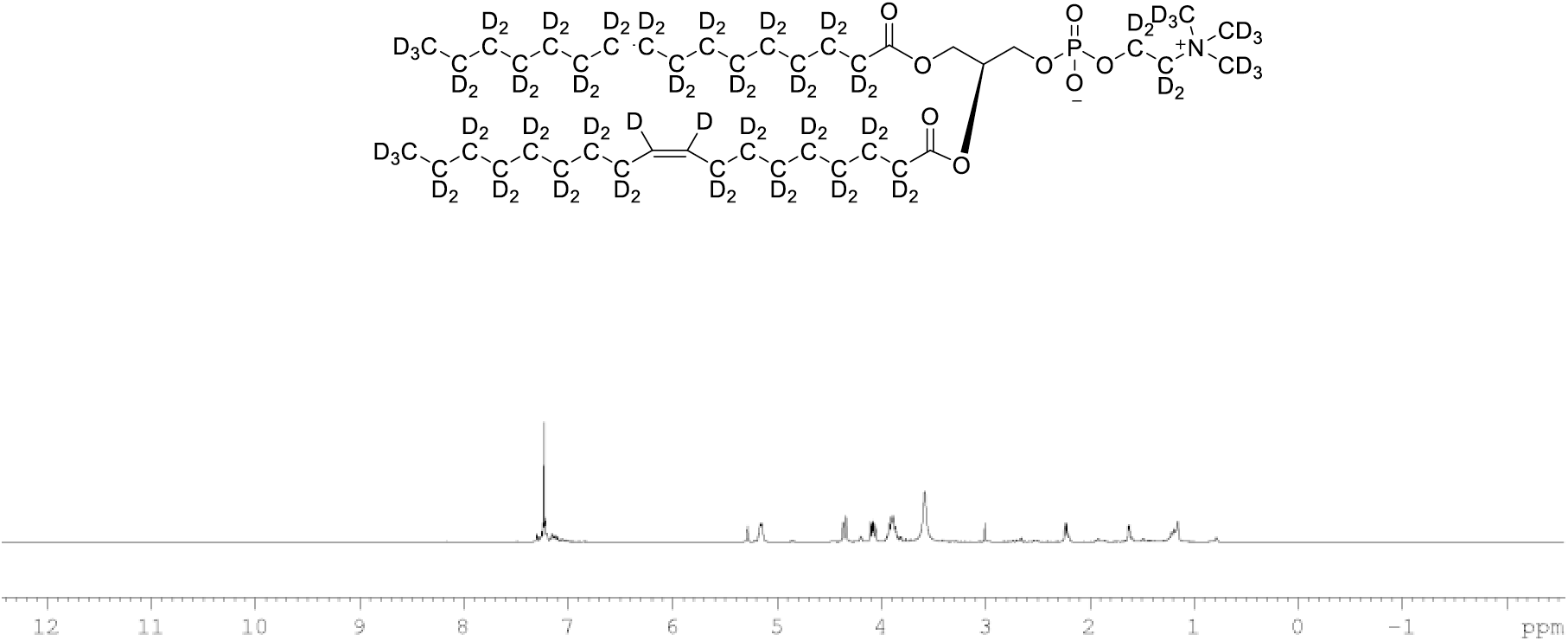
^1^H NMR of POPC-d_77_ in CDCl_3_. (400 MHz, CDCl3), δ residual protons 0.85 (m, 0.17H), 1.25 (m, 1.48H), 1.54 (m, 0.22H), 1.97 (m, 0.18H), 2.22 (m, 0.66H), 3.89 (m, 2H), 4.08 (m, 1H), 4.34 (m, 1H), 5.14 (m, 1H), 5.27 (s, 0.43H)

**Figure supplement 15.**
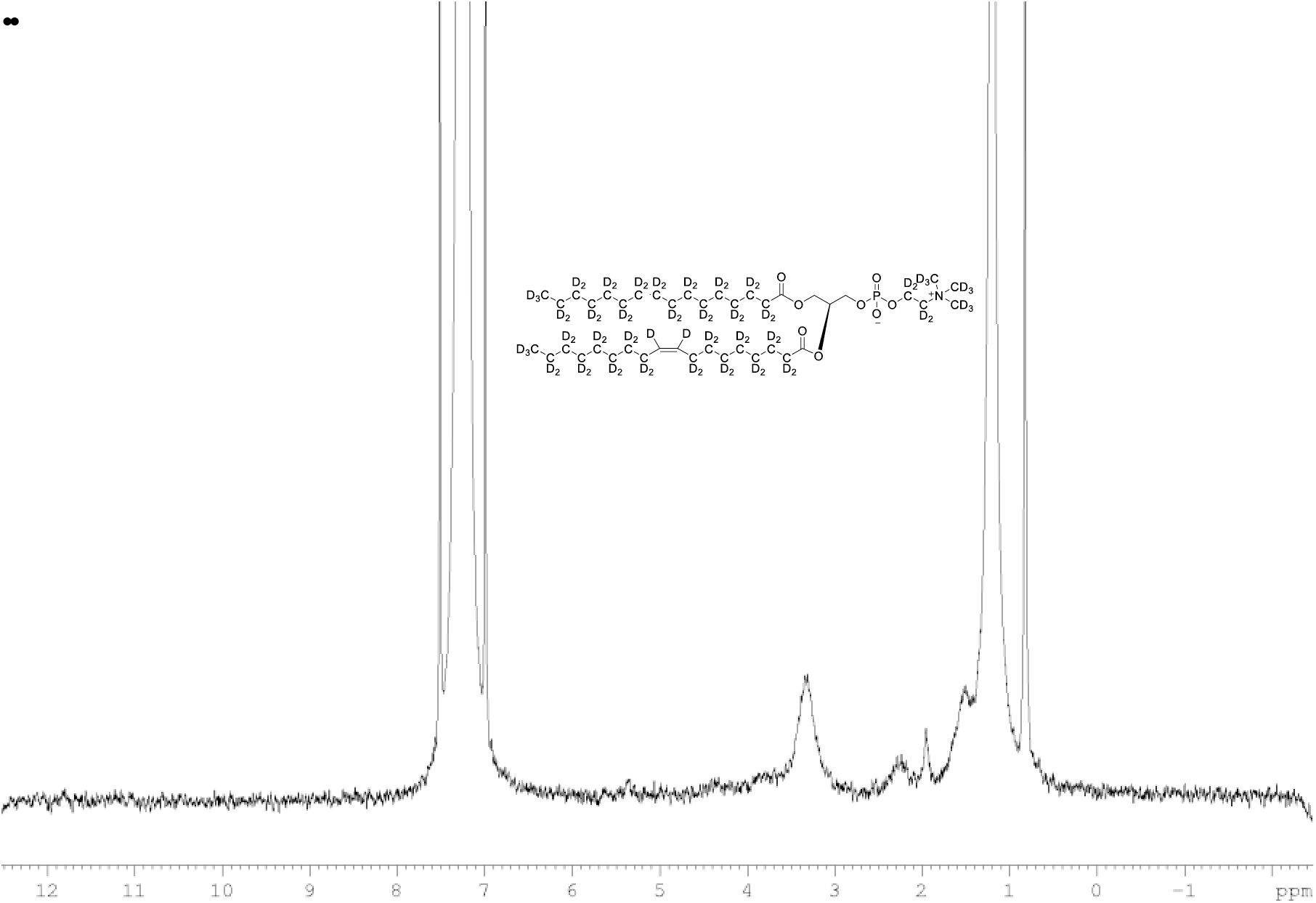
^2^H NMR of POPC-d_77_ in CDCl_3_. (400 MHz, CDCl_3_), δ 0.80 (m), 1.19 (m), 2.20 (m), 1.93 (m, 6.0D), 3.35 (m), 3.84 (m), 5.36 (m).

**Figure supplement 16.**
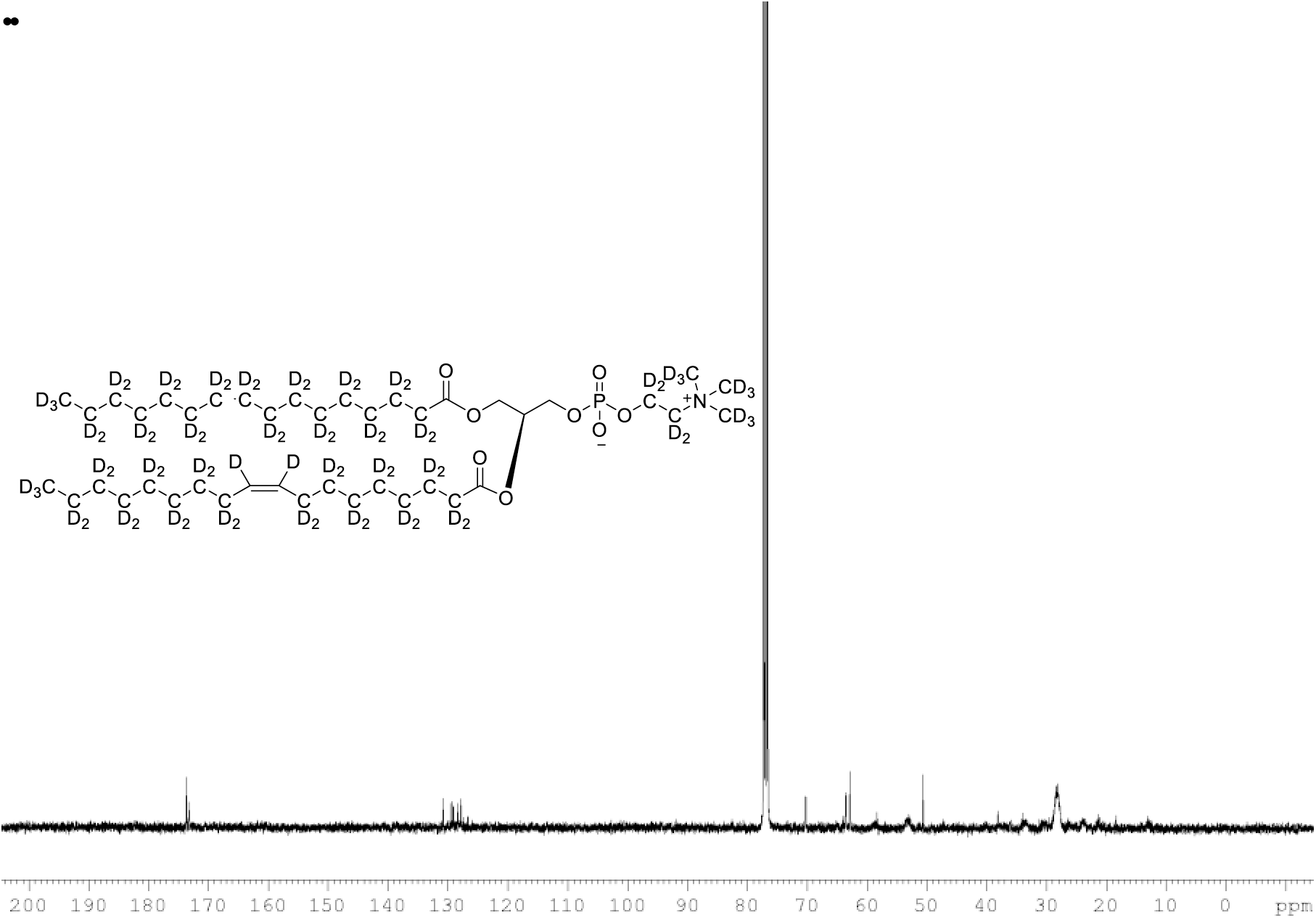
^13^C NMR of POPC-d_77_ in CDCl_3_. (400 MHz, CDCl_3_), δ 13.03 (m), 21.48 (m), 23.88 (m), 26.24 (m), 28.36 (m), 30.49 (m), 33.72 (m), 53.17, 58.76 (m), 62.24 (m), 69.84 (m), 127.90 (s), 129.5, 173.31, 173.69

**Figure supplement 17.**
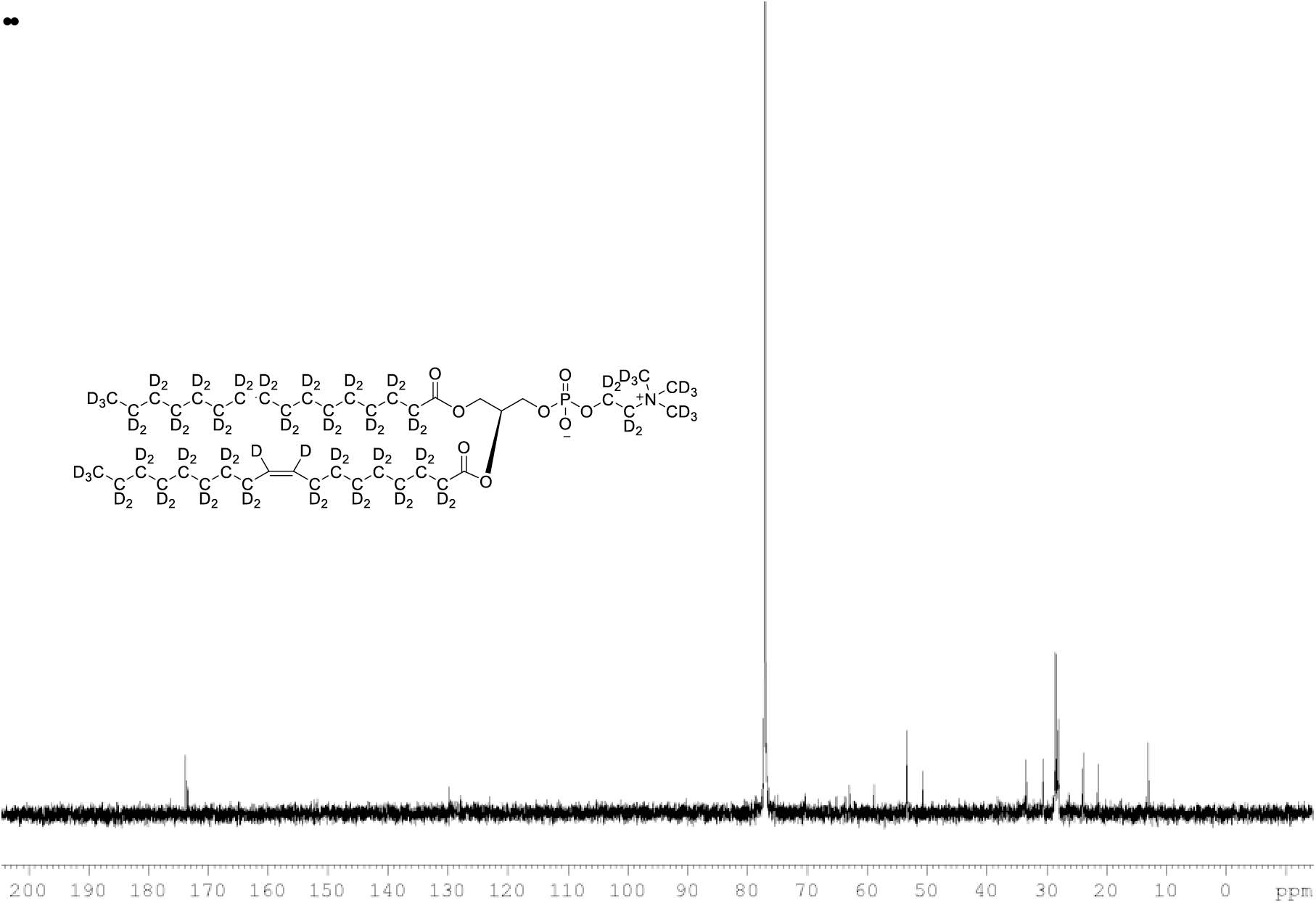
{^1^H}and {^2^H} decoupled ^13^C NMR spectra of POPC-d_77_ in CDCl_3_.

**Figure supplement 18.**
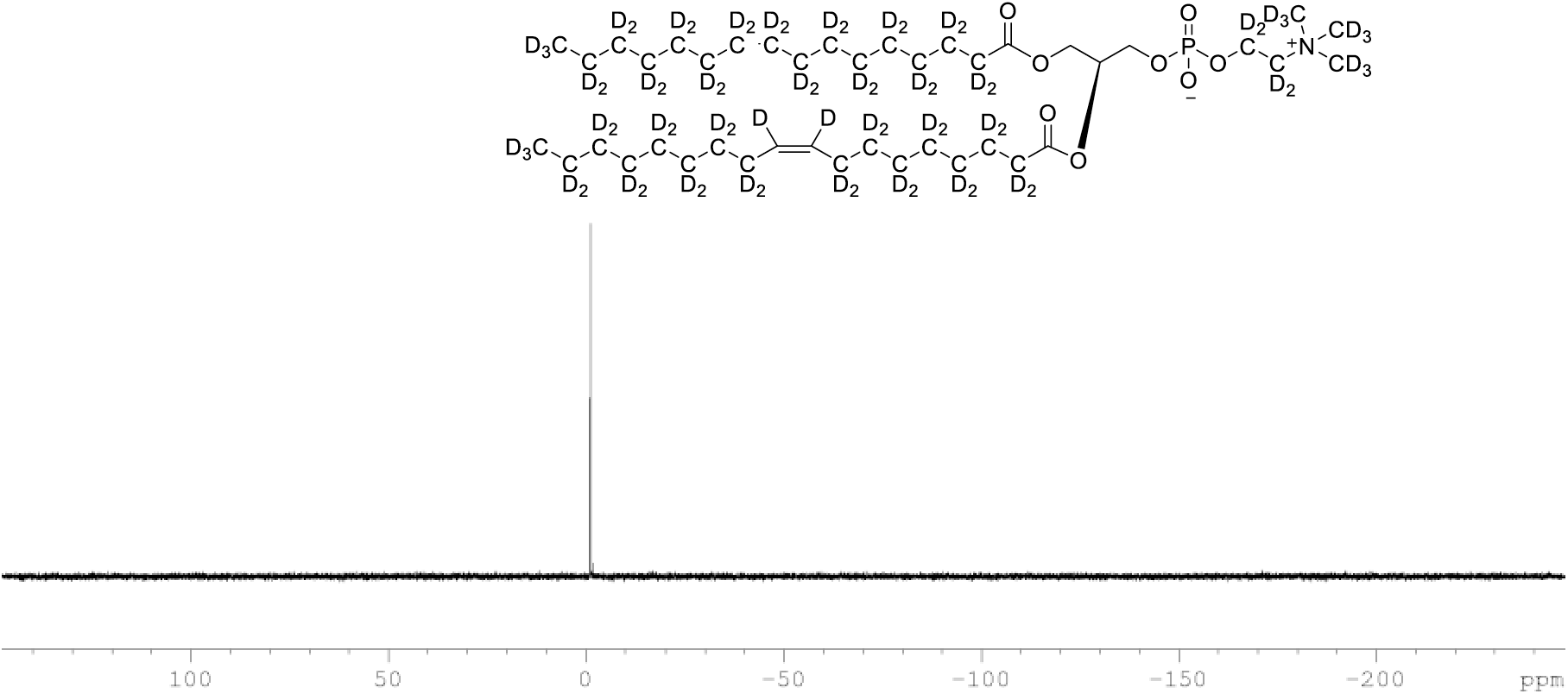
^31^P NMR of POPC-d_77_ in CDCl_3_. (400 MHz, CDCl3), single peak at δ -2.20

**Figure supplement 19.**
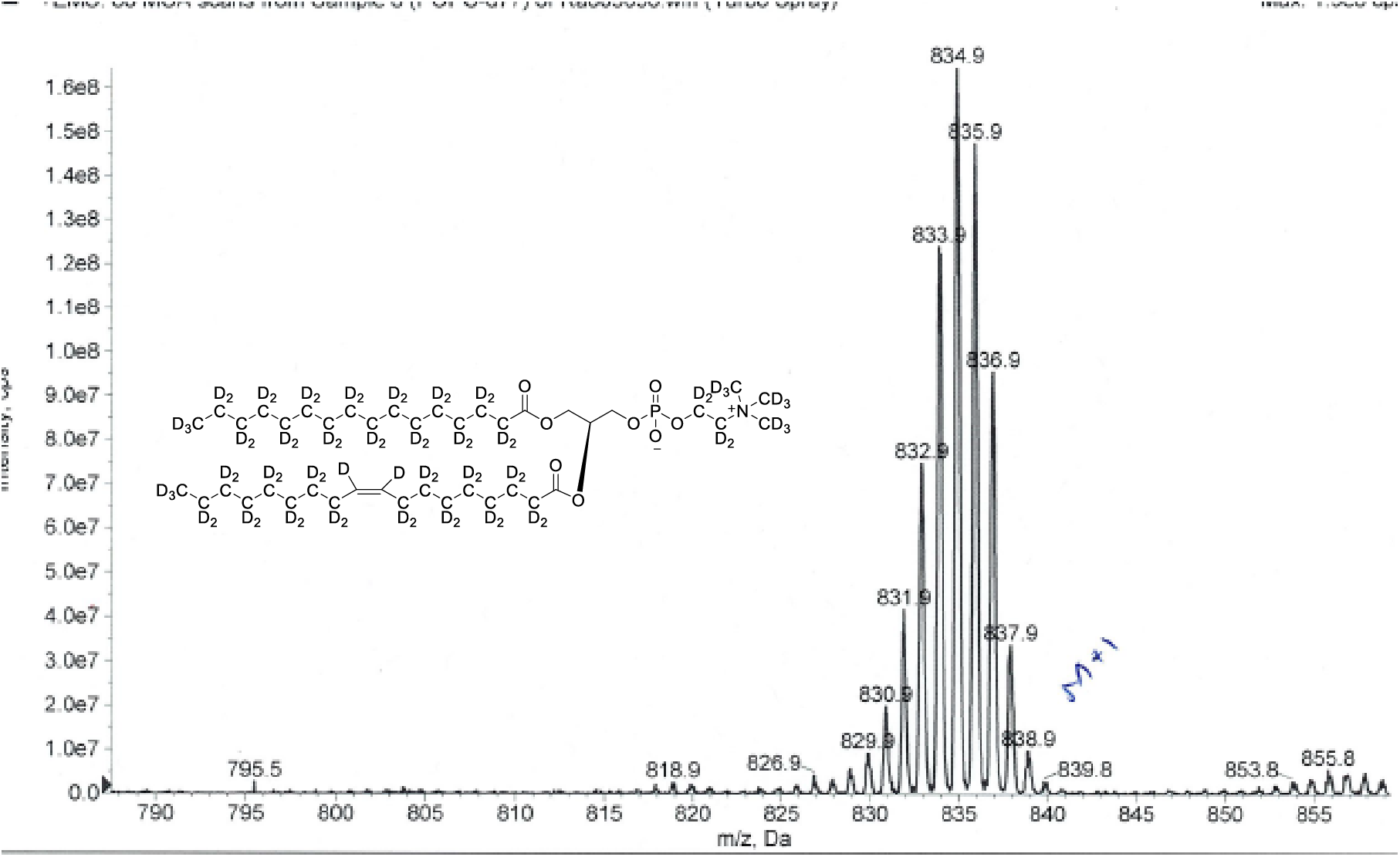
ESI-MS, m/z 838 [M^+1^]^+^ of POPC-d_77_. Overall 93%D, isotope distribution d_77,_ 0%, d_76,_ 0%, d_75,_ 7.5%, d_74,_ 21.2%, d_73,_ 28.7%, d_72,_ 25.3%, d_71,_ 15.5%, d_70,_ 10.8%.

**Figure supplement 20.**
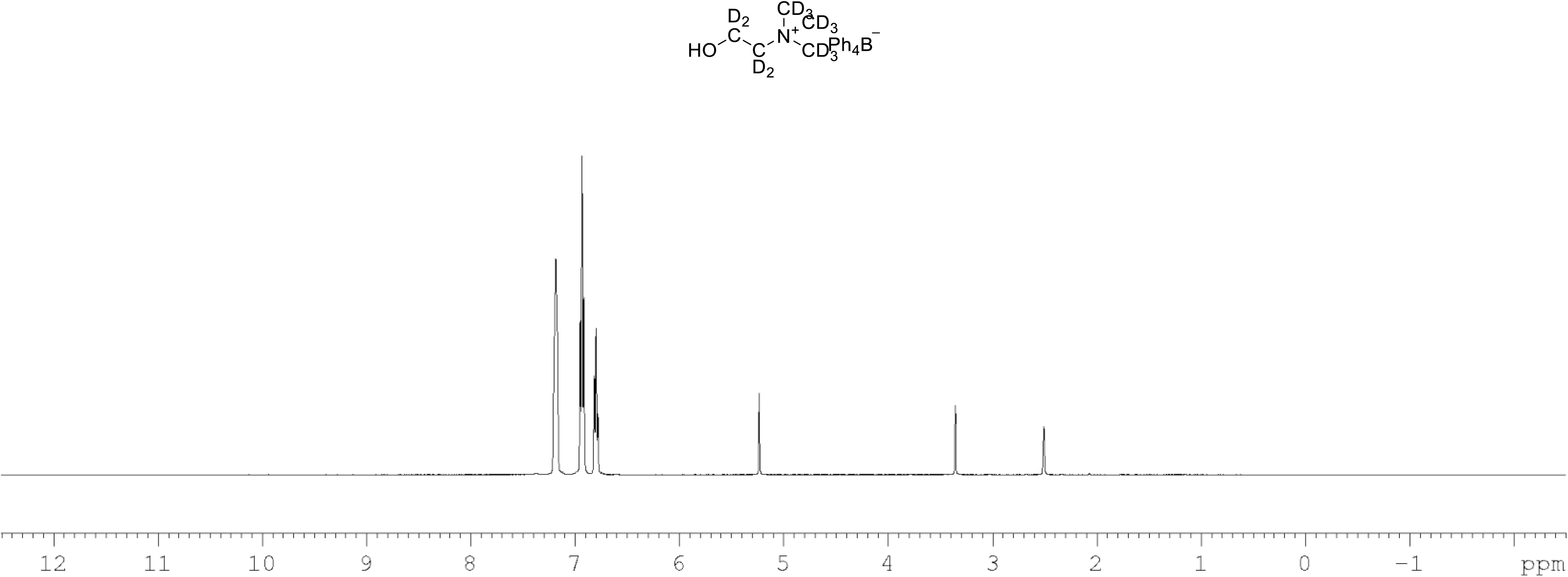
^1^H NMR of choline-d_13_ tetraphylborate (Figure 7, molecule **8**) in DMSO-d_6_. (400 MHz, DMSO-d_6_) δ 6.85 (m, 4H), 6.96 (m, 8H), 7.21 (m, 8H).

**Figure supplement 21.**
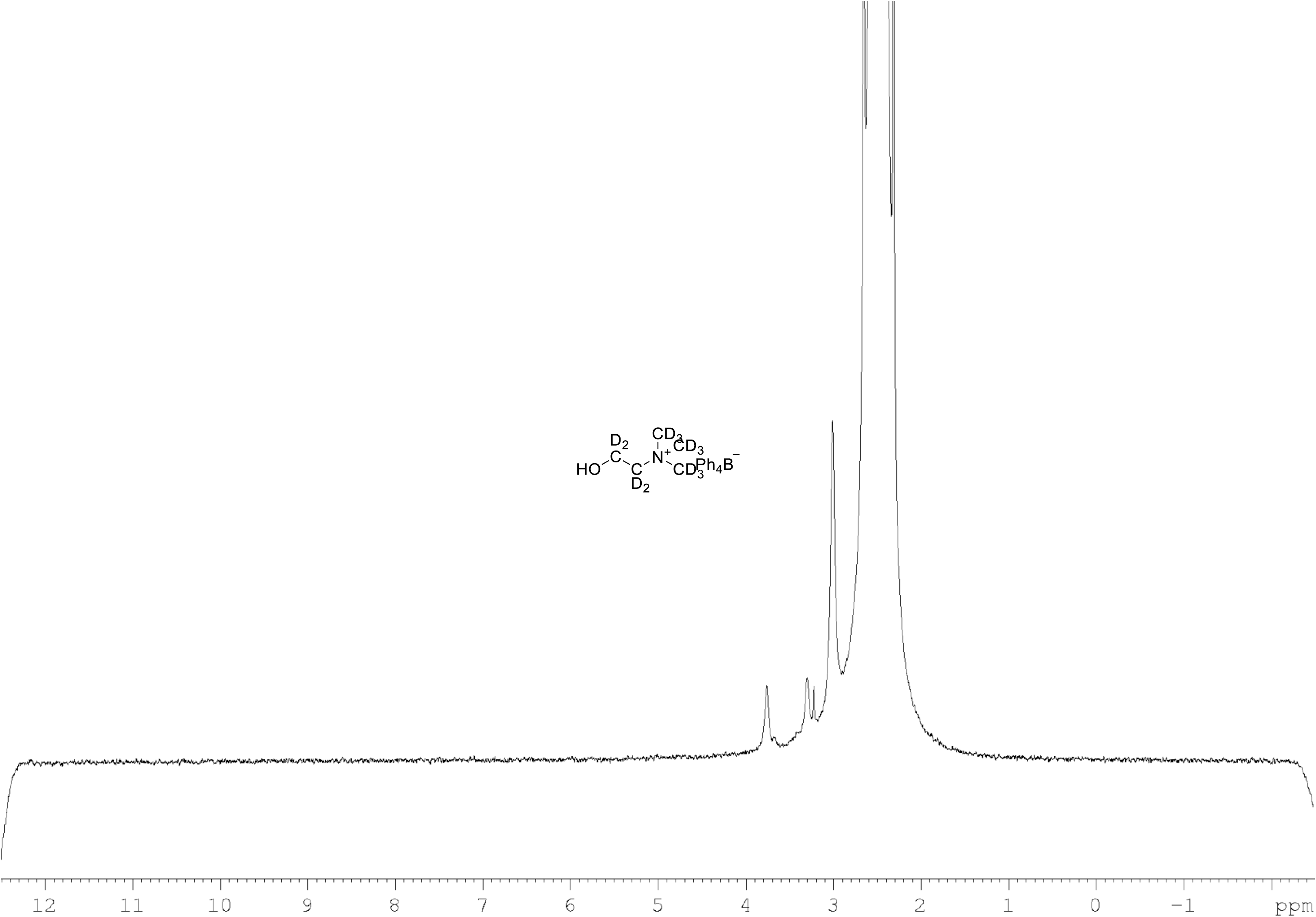
^2^H NMR of choline-d_13_ tetraphylborate (Figure 7, molecule **8**) in DMSO-d_6_. (61.4 MHz, DMSO-d_6_) δ 3.30 (m, 9D), 3.32 (m, 2D), 3.78 (m, 2D).

**Figure supplement 22.**
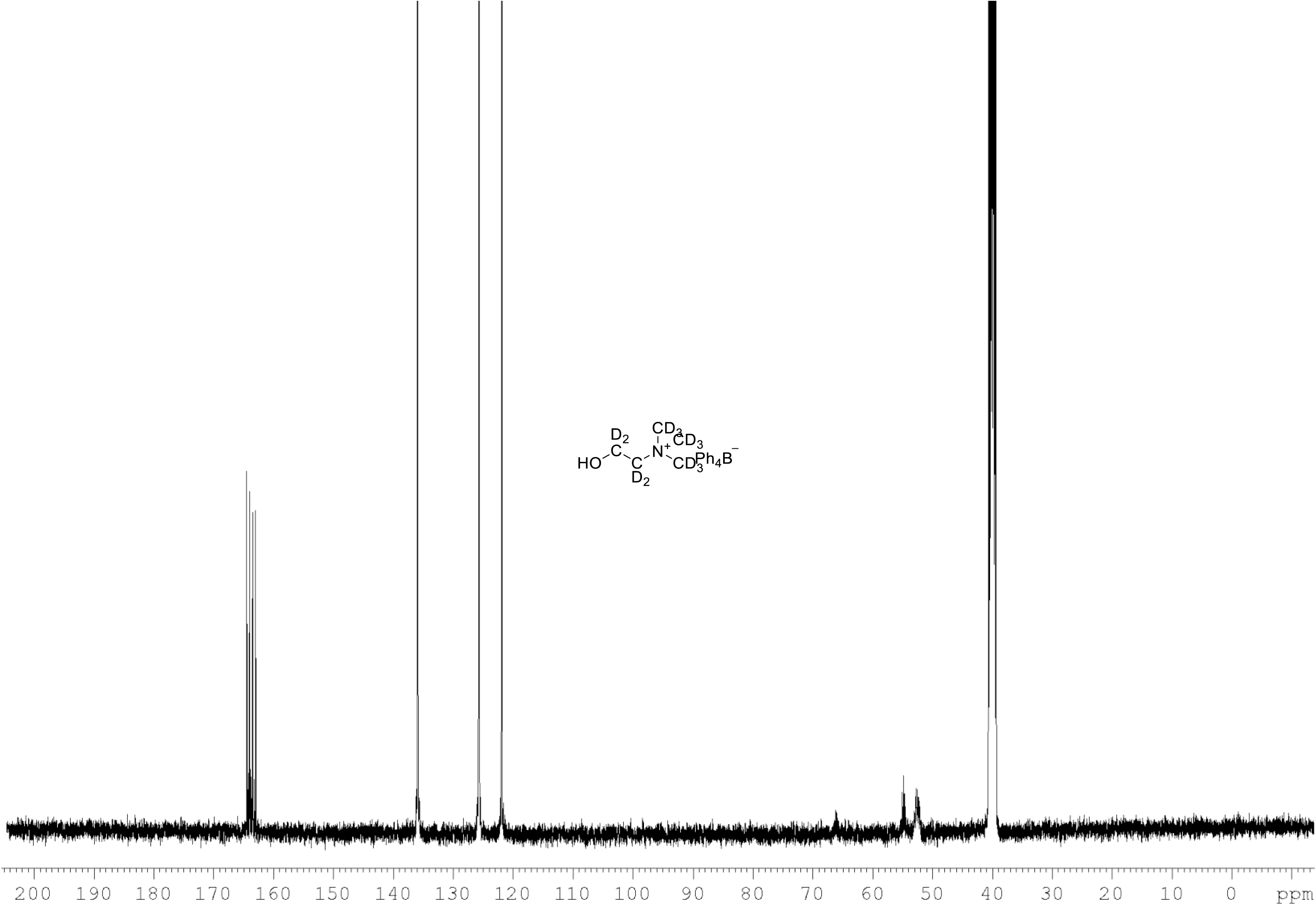
^13^C NMR of choline-d_13_ tetraphylborate (Figure 7, molecule **8**) in DMSO-d_6_. (100 MHz, DMSO-d_6_) δ 52.5 (m), 54.9 (m), 121.9, 125.9, 136.2, 163.9 (m).

**Figure supplement 23.**
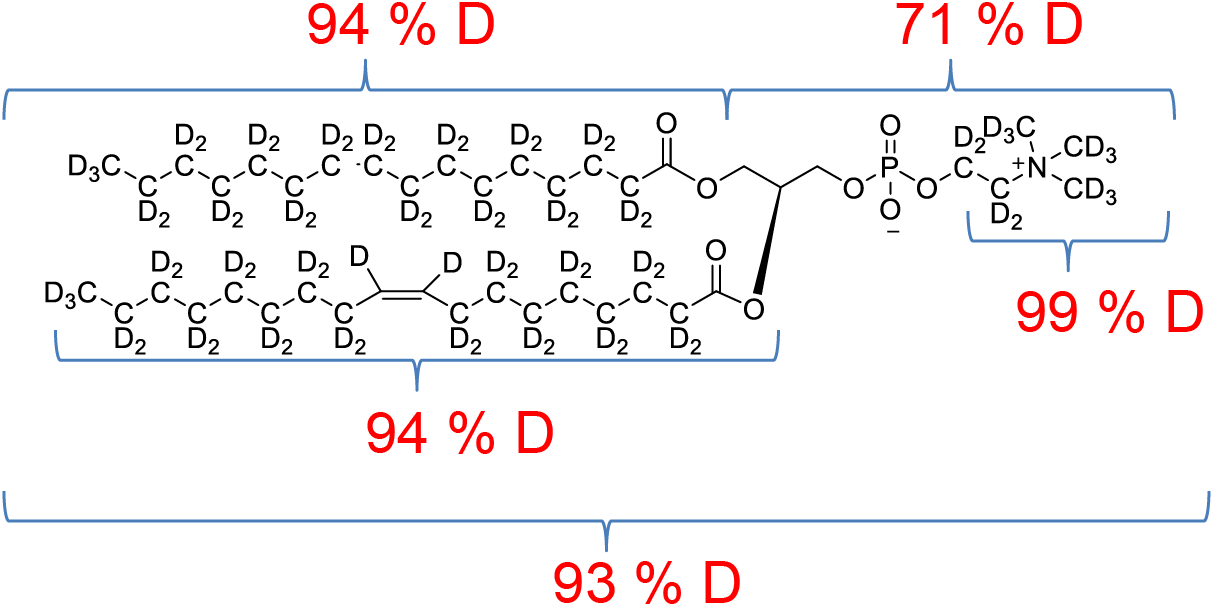
POPC-d_77_ percentage distribution of D levels at different sites.

